# Assessment of structural behaviour of new L-asparaginase and SAXS data-based evidence for catalytic activity in its monomeric form

**DOI:** 10.1101/2023.01.01.522448

**Authors:** Kanti N. Mihooliya, Jitender Nandal, Nidhi Kalidas, Ashish, Subhash Chand, Dipesh K. Verma, Mani S. Bhattacharyya, Debendra K. Sahoo

## Abstract

The present study reports the structural and functional characterization of a new glutaminase-free recombinant L-asparaginase (PrASNase) from Pseudomonas resinovorans IGS-131. PrASNase showed substrate specificity to L-asparagine, and its kinetic parameters, K_m_, V_max_, and k_cat_ were 9.49×10^-3^ M, 25.13 IUmL^-1^min^-1^, and 3.01×10^3^ s^-1^, respectively. The CD spectra showed that PrASNase consists of 30.9% α-helix and 69.1% other structures in its native form. FTIR was used for the functional characterization, and molecular docking predicted that the substrate interacts with serine, alanine, and glutamine in the binding pocket of PrASNase. Different from known asparaginases, structural characterization by small-angle X-ray scattering (SAXS) and analytical ultracentrifugation (AUC) unambiguously revealed PrASNase to exist as a monomer in solution at low temperatures and oligomerized to a higher state with temperature rise. Through SAXS studies and enzyme assay, PrASNase was found to be mostly monomer and catalytically active at 37°C. Furthermore, this glutaminase-free PrASNase showed killing effects against WIL2-S and TF-1.28 cells with IC_50_ of 7.4 µg.mL^-1^ and 5.6 µg.mL^-1^, respectively. This is probably the first report with significant findings of fully active L-asparaginase in monomeric form using SAXS and AUC and demonstrates the potential of PrASNase in inhibiting cancerous cells, making it a potential therapeutic candidate.

**HIGHLIGHTS:**

- A new L-asparaginase (PrASNase) was structurally and functionally characterized.
- SAXS revealed PrASNase is functionally active in monomeric form and oligomerizes with temperature rise.
- Monomeric PrASNase showed an IC_50_ value of 7.4 and 5.6 µg mL^-1^ against WIL2-S and TF-1.28 cells.
- Cytotoxicity of PrASNase against leukemic cell lines showed its potential as a biotherapeutic.

**GRAPHICAL ABSTRACT:** 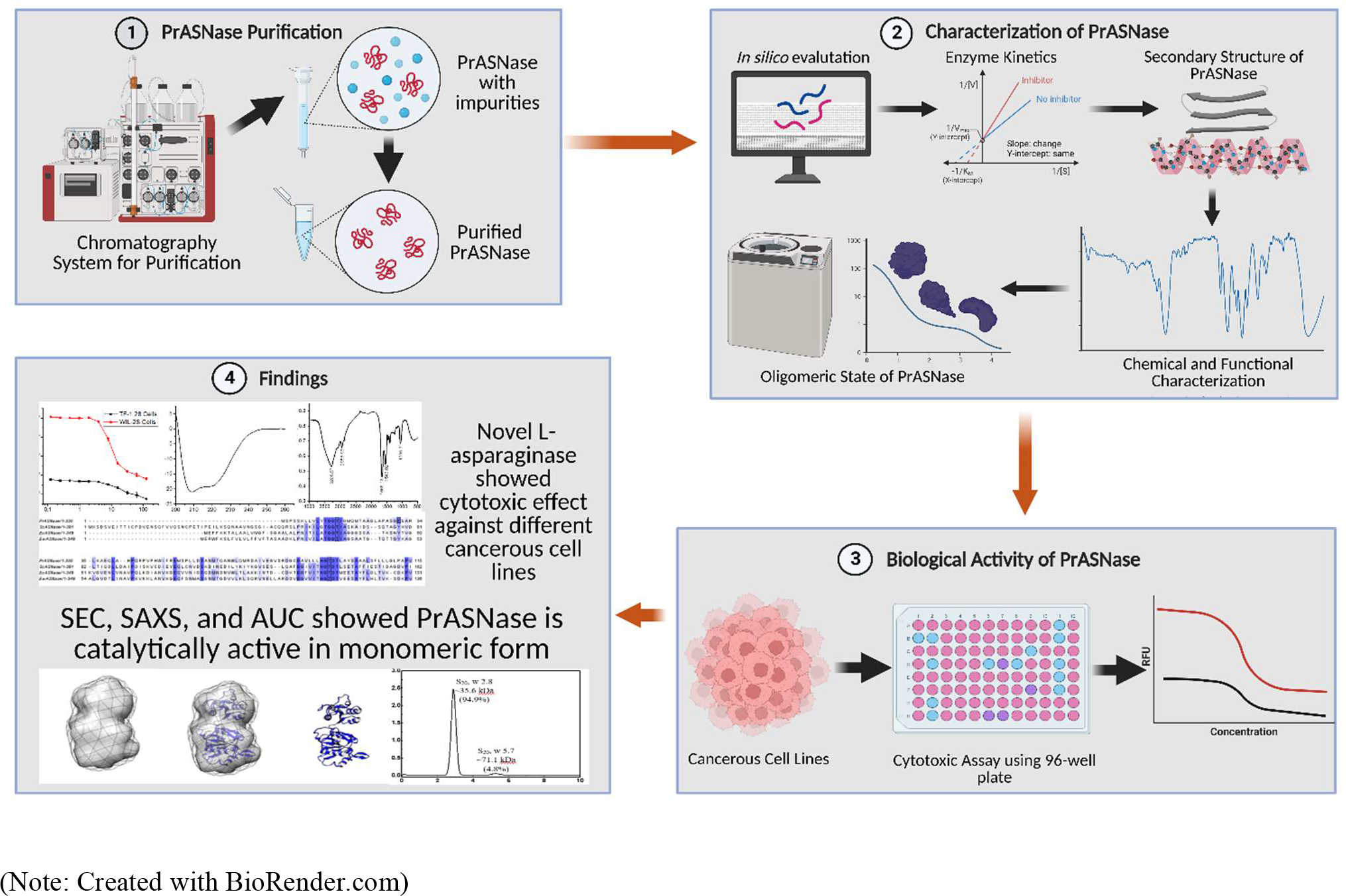

## 1. Introduction

L-Asparaginase (EC 3.5.1.1) is an important enzyme having diverse applications in the pharmaceutical and food industries as a potential agent for the treatment of acute lymphoblastic leukaemia (ALL) and inhibition of acrylamide formation in baked and fried foods [1, 2]. The mechanism of L-asparaginase to hydrolyze L-asparagine into aspartic acid and ammonia and the lack of ability of leukemic cells to synthesize its L-asparagine make L-asparaginase an effective therapeutic for the treatment of ALL [3–5]. This enzyme hydrolyzes bloodstream L-asparagine into aspartic acid and ammonia and creates an L-asparagine-deprived environment, which results in the inhibition of leukemic cells and several other human cancerous cells [4,6–8]. Furthermore, L-asparaginase has another essential application in the inhibition of the formation of acrylamide from fried and baked carbohydrate-based food products, as the presence of higher levels of acrylamide in these food products has raised a worldwide health concern [9, 10].

Commercially, L-asparaginase from Escherichia coli and Erwinia chrysanthemi is used for the treatment of ALL [11]. There are several forms of L-asparaginases available in the market, including native L-asparaginases (Elspar® [Merck, 2000] from E. coli and crisantaspase [Medicines Evaluation Board, 2000] from Erwinia chrysanthemi), pegylated L-asparaginase (pegaspargase, Oncaspar® [European Medicines Agency, 2016] from E. coli), and the recombinant L-asparaginase (Spectrila® [European Medicines Agency, 2015] expressed in E. coli as a host) [5,12,13]. Furthermore, L-asparaginases from these sources are immunogenic and have high glutaminase activity, which results in severe side effects during the treatment [3,12,14–16]. In addition, L-asparaginase from L-Aspergillus oryzae and Aspergillus niger are currently used under the brand name of Acrylaway® from Novozymes A/S (Bagsvaerd,

Denmark) and PreventAse™ from DSM (Heerlen, The Netherlands), respectively, in the food industry for mitigation of acrylamide [5,10,13].

A protein’s structural and functional characterization for understanding stability and oligomeric conditions is essential for commercial exploitation. The L-asparaginase activity is known to depend on its source and multimeric state [17]. For example, the commercial L-asparaginase (i.e., Kidrolase) from E. coli was natively tetramer, but upon storage at -80°C, its oligomeric state was reported to change to monomer and another higher-order state which was less active as compared to tetramers [18]. Another report showed that L-asparaginase from Pyrococcus furiosus was natively dimer and fully functional, whereas, under several conditions, it converted to a monomer that exhibited low or no activity [19]. Several studies have reported that L-asparaginase from different sources originated in various oligomeric states like dimeric, hexameric, and tetrameric [3,17,20–22].

The current study focused on the structural and functional characterization of new glutaminase-free recombinant L-asparaginase from Pseudomonas resinovorans IGS-131 (PrASNase). The protein was purified and assessed for conserved motifs and binding pockets using multiple sequence alignment and molecular docking, respectively. The chemical and functional groups were analyzed by application of Fourier transform infrared spectroscopy (FTIR). The secondary structure of PrASNase was determined by circular dichroism (CD). The substrate specificity of the purified PrASNase was evaluated using different substrates, and the Michaelis-Menten equation was used to determine its kinetic parameters. The structural characterization of PrASNase was carried out using SEC, SAXS, and AUC techniques, and the anti-cancerous potential of the purified enzyme was checked against WIL2-S and TF-1.28 cell lines.

## 2. Materials and Methods

### 2.1. Chemicals

Nessler’s reagent, L-asparagine, trichloroacetic acid (TCA), D-asparagine, L-glutamine, potassium bromide (KBr), D-glutamine, L-aspartic acid, D-aspartic acid, RPMI-1640, Luria Bertani (LB) broth, fetal bovine serum (FBS), glucose, penicillin, streptomycin, and sodium pyruvate were purchased from Himedia, India. Chloramphenicol and kanamycin were bought from MP Biomedicals, LLC, USA. Isopropyl-β-D-1-thiogalactopyranoside (IPTG) was purchased from UBPBio, USA. Reagents and protein markers used in sodium dodecyl sulfate-polyacrylamide gel electrophoresis (SDS-PAGE) were obtained from BioRad, India. Resazurin sodium salt (Alamar Blue) was purchased from Sigma, USA. All other chemicals used in this study are of analytical grade.

### 2.2. Purification of recombinant L-asparaginase (PrASNase)

The cells were harvested from the fermented broth by centrifugation. The lysis of harvested cells by sonication was followed by separating and loading the periplasmic fraction onto the Ni-NTA column. The bound protein was then eluted using imidazole, and the remaining impurities were removed by gel filtration chromatography as described before [23].

### 2.3. Assay of asparaginase activity

L-asparaginase activity was quantified using the Nesslerization method to determine ammonia, as described by Imada et al. and Mihooliya et al. [23, 24]. The enzyme (L-asparaginase) activity was expressed in the international unit (IU), one IU of the enzyme being referred to as the quantity of enzyme required to release 1 µmole of ammonia per min per ml at pH 8.6 and 37°C [1,24,25].

### 2.4. Sequence alignment and molecular docking of PrASNase gene

The nucleotide sequence encoding for the PrASNase gene obtained in the present study (GenBank accession number MK799854) was translated into the amino acid sequence using the translation tool at the ExPASy online server (https://web.expasy.org/translate/) [26]. The amino acid sequence of PrASNase was compared with the sequence of E. coli asparaginase (EcASNase, Uniprot number P00805), Erwinia chrysanthemi asparaginase (EwASNase, Uniprot number P06608), and Saccharomyces cerevisiae asparaginase (ScASnase, Uniprot number P38986) available in the database using protein data bank (PDB) Blast (https://www.rcsb.org/). The amino acid sequence was analyzed and aligned using Clustal Omega (https://www.ebi.ac.uk/Tools/msa/clustalo/) and Jalview [26–28].

The three-dimensional structure of PrASNase was prepared using an I-TASSER server [29] that provides automated tools for protein structure prediction and analysis [30]. The monomeric structure of PrASNase was further used for docking studies with L-asparagine substrate using HADDOCK 2.4 web server. We followed all standard HADDOCK parameters and constraints for docking, which generated 1000 predicted models in the rigid body minimization stage. Out of 1000 best, 200 models were further refined based on energy function. The final models are grouped based on a specific similarity measure, such as positional interface ligand RMSD or the fractions of common contacts. The RMSD value from the overall lowest-energy structure is 0.6, and the Z score is -0.1.

### 2.5. Physicochemical characterization of purified PrAsnase

#### 2.5.1. Molecular weight determination

The molecular weight profile of PrASNase was studied by SEC-HPLC using a gel filtration column (Yarra® SEC-3000, Phenomenex, USA) connected to the HPLC system (Shimadzu Corporation, Japan). The column was equilibrated with 20 mM Tris-Cl (200 mM NaCl and pH 8.0) and operated at 25°C and 1.0 mL min^-1^ of flow rate. In our previous study, gel filtration chromatography with Superdex-75 SEC column (Hi-Load 16/600, Prep Grade, GE Healthcare, USA) and MALDI analysis were used to determine the molecular mass of PrASNase [23].

#### 2.5.2. Substrate specificity

Different substrates (such as L-asparagine, L-aspartic acid, L-glutamine, D-asparagine, D-glutamine, and D-aspartic acid) were used to evaluate L-asparaginase activity. The reaction mixture consisted of each of the above-said substrates at a concentration of 10 mM and L-asparaginase at a final concentration of 500 nM under assay conditions. Results obtained were represented in percentage relative activity.

#### 2.5.3. Kinetic parameters

The kinetic parameters, i.e., Michaelis-Menten constant (K_m_), maximal velocity (V_max_), and turnover number (k_cat_) of purified PrASNase, were determined by nonlinear regression of Lineweaver-Burk plots [31]. Under assay conditions, the varying concentrations of substrate L-asparagine (2-20 mM) and a constant enzyme (protein) concentration (500 nM) were used.

#### 2.5.4. Circular dichroism (CD) spectroscopy

The secondary structure of PrASNase was determined by monitoring its CD spectra with a Jasco J-815 spectropolarimeter, USA. The CD spectra of protein (0.25 mg mL^-1^) contained in a quartz cuvette of 1 mm path length were recorded between 200 to 260 regions at 20°C [19]. Three accumulations of each scan were recorded, averaged, and subtracted from baseline 20 mM Tris-Cl (200 mM NaCl and pH 8.0).

#### 2.5.5. Fourier transform infrared spectroscopy (FTIR)

FTIR method was used to determine the chemical bonds and functional groups present in the purified PrASNase. The lyophilized protein sample of 0.5 mg concentration was mixed with 100 mg KBr for FTIR analysis. The sample spectrum was recorded and analyzed on the FTIR spectrometer (Vertex 70, Bruker Optik GmbH, Germany) between 4000-400 wavenumbers per cm.

#### 2.5.6 Oligomeric state analysis using size exclusion chromatography (SEC)

The changes in the oligomeric state of PrASNase were studied using the gel filtration column Superdex-200 (10/300 GL Increase, GE Healthcare, USA). The PrASNase was freshly purified and distributed equally in two different vials. In one vial, the native state of PrASNase was arrested using the crosslinking agent glutaraldehyde, and the other vial was left without the glutaraldehyde. Both vials were stored at 4°C, and the oligomeric state was analyzed using the SEC at 0 h, 24 h, and 48 h. The protein molecular weight standards profile was also plotted using the same SEC column for comparison (Fig. 6G).

#### 2.5.7. Small Angle X-ray Scattering (SAXS)

All SAXS datasets mentioned in this paper were obtained at the CSIR-Institute of Microbial Technology using in-house SAXS equipment (SAXSpace, Anton Paar GmbH, Graz, Austria). Sealed X-ray tube generated line collimated X-rays that were incident for 1 h on the PrASNase sample (and matched buffer) filled in a 1 mm diameter capillary (thermostated quartz). The data was collected for L-asparaginase solution from 10°C to 70°C (283 to 343 K) and backward at an interval of 10°C [32, 33]. A 1D CMOS Mythen detector was used to collect the scattered data (Dectris, Switzerland). Supplementary Table S2 lists all the programs used to process the obtained data. Using the SAXSTreat program, the beam location was corrected. The beam profile was exploited for data desmearing to represent point collimation using the SAXSQuant program after the subtraction of buffer contribution. Finally, the data was acquired as scattering intensity I(q) as a function of q, where q is the momentum transfer vector and is defined as q = 4π (sinθ)/λ with θ being the scattering angle and λ is the wavelength of X-rays used in nm. The ATSAS suite of programs version 3.0 was used to process the datasets to obtain the radius of the gyration of the molecule (R_g_). The normalized Kratky analyses were carried out in an automated mode, and the plots were manually plotted for the generation of the figure. The scattering shapes of the molecule were reconstructed ab initio using a chain-like ensemble of dummy residues equal to the number of amino acids in the protein. Ten models were computed, aligned by inertial axes, and averaged to obtain the envelope maps of L-asparaginase in the solution state. All the plots and models were made using Origin Lab software version 5 and UCSF Chimera version 1.14.

Additionally, the Swiss Modeller Server was utilized to search for sequence similarity-based structural template selection and generation of the representative model of PrASNase. ProMod3 version 2.0.0 was used to select one chain from PDB ID 3NTX solved at a resolution of 1.9 Å (Cytoplasmic L-asparaginase I from Yersinia pestis). The program suggested a homodimer and a sequence similarity of only 35.69. Sequence coverage was high at 0.98, with some gaps and additional segments in the target vs template sequence. The final energy-optimized model of PrASNase was provided with a global model quality estimation (GMQE) of 0.73. Later, this model matched well with the SAXS data-based structural parameters and the restored shape model (Supplementary Table S1) [32, 33].

#### 2.5.8. Sedimentation velocity analytical ultracentrifugation (SV-AUC)

Sedimentation velocity analytical ultra-centrifugation (SV-AUC) was carried out to analyze the presence of different oligomer species in the purified PrASNase using Beckman Coulter XL-1 AUC (California, USA). PrASNase at a concentration of 0.5-0.8 mg mL^-1^ was taken in the sample cell of the AUC cuvette, and 20 mM Tris-Cl (200 mM NaCl and pH 8.0) was used in the reference cell of the cuvette. The rotor speed of the instrument was set at 40000 rpm, and 100 scans were recorded by giving 3 min intervals between each scan. The scans’ monitoring started once the instrument’s temperature reached 20°C. The whole process was performed between 25°C-30°C, and the results of SV-AUC were analyzed using SEDFIT software.

### 2.6. Biological activity of purified PrASNase against cancerous cell lines

#### 2.6.1. Cell lines and culture conditions

The anti-cancerous effect of purified PrASNase was checked on human tumor cells, i.e., WIL2-S (Human spleen B lymphoblast, ATCC CRL-8885) and TF-1.28 (Human bone marrow erythroblast, ATCC CRL-2003). Both cell lines were suspended in RPMI-1640 growth medium supplemented with 10% FBS and antibiotics (penicillin and streptomycin), followed by incubation at 37°C and 5% CO_2_. The cell viability (at least 90%) and the cell count before seeding were carried out using the Automated Cell Counter (Biorad, USA).

#### 2.6.2. Cell-based inhibition method

The antiproliferative activity of PrASNase was checked by the cell-based inhibition method using Alamar blue dye. The protein (PrASNase) concentration of 250 µg mL^-1^ was diluted two-fold to form 11 dilutions of 100 µL in standard 96-well microplates. All dilutions were prepared in duplicates. 100 µL of WIL2-S cells (0.5×10^6^ cells mL^-1^) and TF-1.28 cells (0.5×10^5^ cells mL^-1^) were added separately to these dilutions, and the microplates were incubated for 24 h at 37°Cand 5% CO_2_. Then 50 µL of Alamar blue dye was added to each well, and the plates were further incubated for 24 h at 37°C and 5% CO_2_. After incubation, the plate’s relative fluorescence units (RFU) were read at 530 nm excitation and 590 nm emission using Spectrofluorometer (SpectraMax Gemini Spectrofluorometer, USA).

## 3. Results and discussions

### 3.1. Sequence alignment of PrASNase with other bacterial asparaginases

The protein sequence of PrASNase was aligned with PASNase (Pseudomonas asparaginase), EcASNase (E. coli asparaginase), EwASNase (Erwinia asparaginase), and BsASNase (Bacillus asparaginase) for checking the conserved residues (Fig. 1). The essential amino acids which perhaps involve in various roles during catalysis of the enzyme were primarily found conserved in PrASNase with minor substitutions [28]. First, the tyrosine (Y) of EcASNase and EwASNase were replaced with phenylalanine (F) in PrASNase. The replacement of tyrosine with phenylalanine may not have much effect as both amino acids are aromatic. At the fourth conserved amino acid position, serine (S) in PrASNase, is conserved in Pseudomonas sp. and Bacillus subtilis L-asparaginase. The deduced amino acid sequence of the PrASNase has shown the highest sequence similarity with EcASNase (21.8%) and EwASNase (21.2%). PrASNase has 19.3% and 15.4% sequence similarity with Pseudomonas sp. and Bacillus subtilis asparaginase, respectively.

**Fig. 1.**
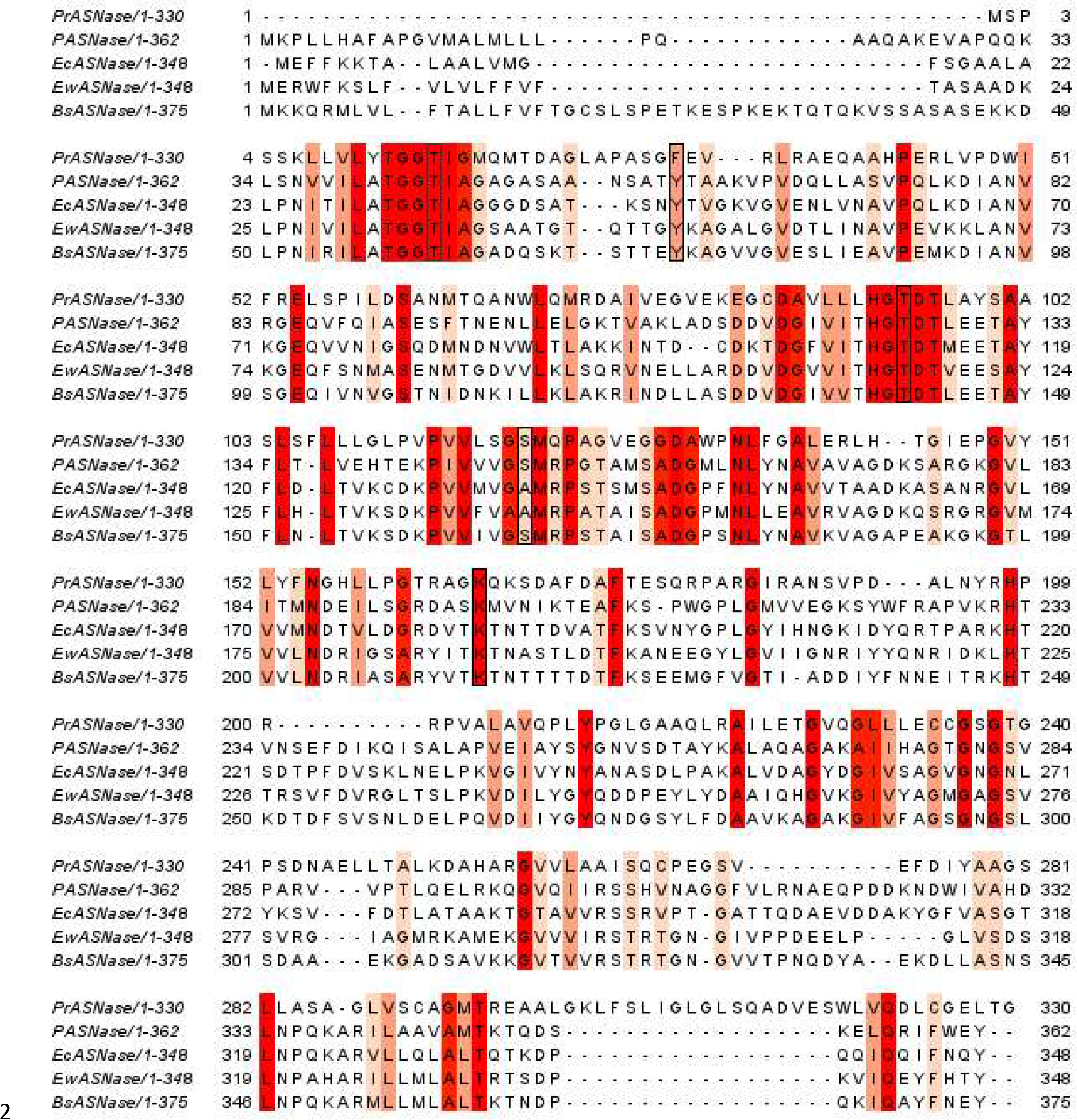
The alignment of PrASNase with EcASNase, EwASNase, BsASNase, and PASNase. The amino acids, which are shaded in red color, represented the percentage identity and conservation between all L-asparaginases with a 50% threshold. The black boxes represented the conserved motifs.

### 3.2. Homology modeling and molecular docking of PrASNase with L-asparagine

Molecular docking showed that L-asparagine binds at the cavity between N- and C-terminal domains of PrASNase (Fig. 2A). The previously solved structures of another asparaginase reveals a conserved binding site for L-asparagine that is located between the N- and C-terminal of the protein. We performed docking of PrASNase with L-asparagine using HADDOCK 2.4. For the PrASNase-L-asparagine complex, HADDOCK clustered 20 structures in the top 4 clusters. These clusters were then manually analysed. The topmost cluster was observed to be the best fit with the HADDOCK score for PilB-PilC, which was 590.2 +/- 23.9 and a Z-score of -0.1. The docking result obtained for this structure has a cluster size of 11 and has Van der Waals energy of -11.3 +/- 3.6, Electrostatic energy of -29.3 +/- 8.8 and buried surface area of 324.4 +/- 2.9. The molecular docking results showed that the L-asparagine fits in the conserved cavity of PrASNase formed between the N- and C-terminal domain (Fig. 2B). The hydrophilic residues that surround this predicted binding cavity allow L-asparagine to directly interact with alanine 98, serine 103, and glutamine 66 through hydrogen bonds (Fig. 2C). Docking study of PrASNase with L-asparagine highlights that L-asparagine binds at the same conserve binding pockets that were previously discovered in both E. coli and Bacillus species asparaginases [34, 35]. This will also imply that other asparaginases and PrASNase have a significant common mechanism of action. The interesting finding is that serine has been substituted for threonine in the active site. This could account for its involvement in relatively high activity and capturing substrate in monomeric form.

**Fig. 2.**
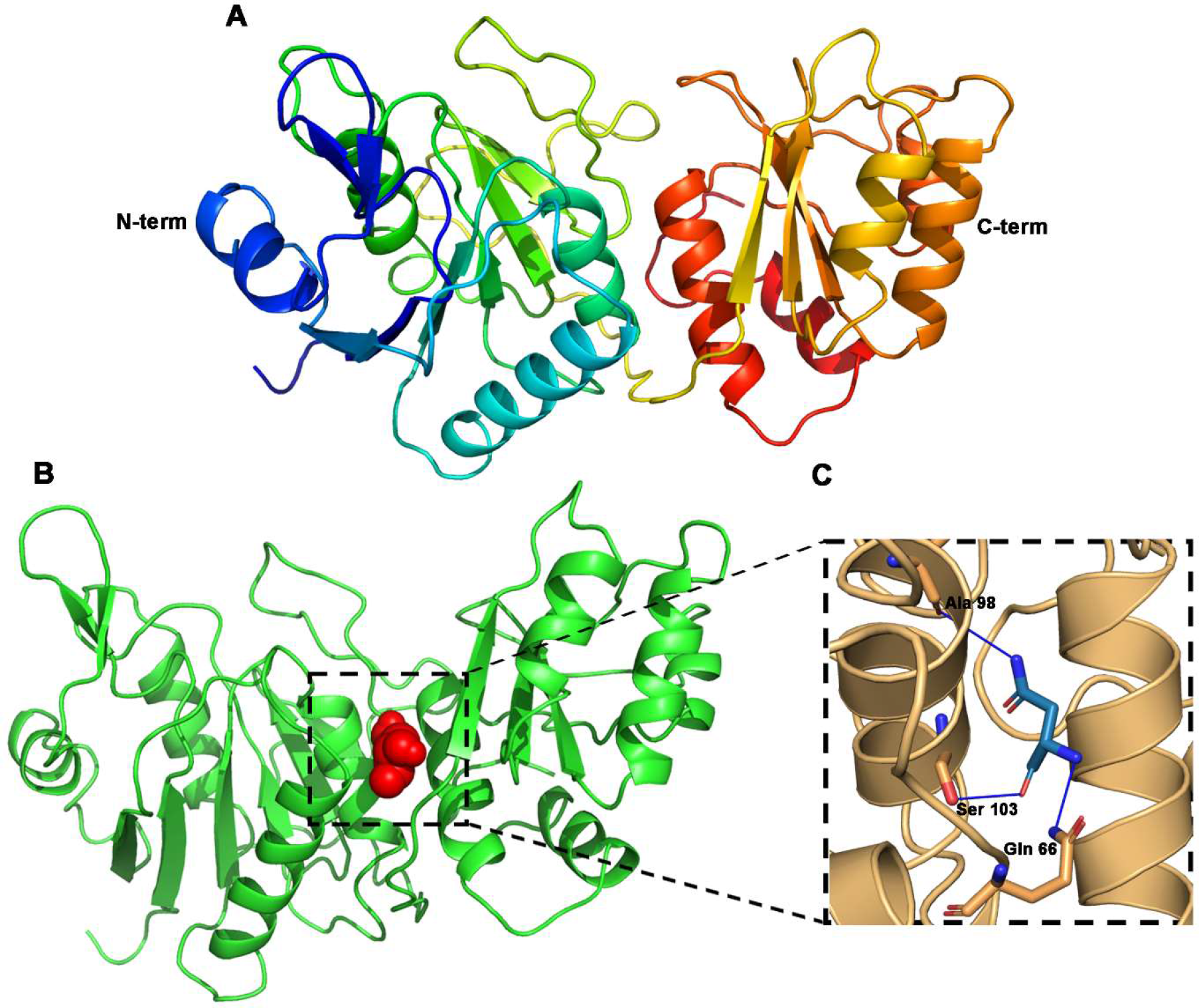
Molecular docking studies of PrASNase with L-asparagine (A) Predicted three-dimensional models of PrASNase show its N and C-terminal domain connected with a linker residue (B), and (C) Schematic showing the active site and residues of the PrASNase that interact with L-asparagine.

### 3.3. Molecular mass determination

The SEC-HPLC analysis of the purified PrASNase showed an individual peak at a retention time of 9.106 min at 280 nm, corresponding to the molecular weight of approximately 35 kDa (Fig. 3A). The peak was collected and run on 12% SDS-PAGE gel under reducing conditions. A prominent monomeric protein band around 35.6 kDa was observed (Fig.3B).

**Fig. 3.**
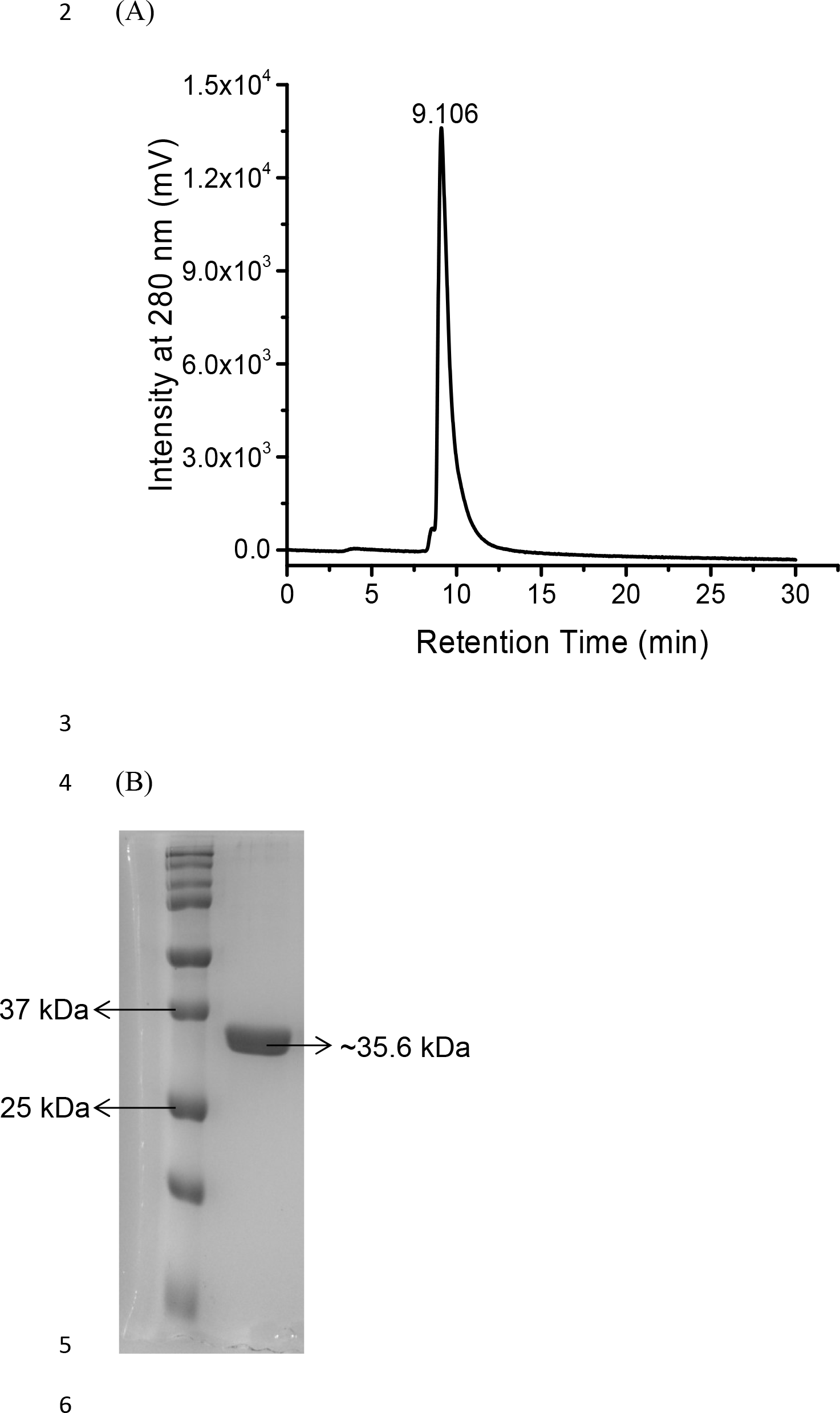
Molecular mass confirmation of PrASNase (A) SEC-HPLC chromatogram of the purified PrASNase, and (B) SDS-PAGE analysis of purified PrASNase.

### 3.4. Substrate specificity and kinetic parameters determination

The substrate specificity of PrASNase was evaluated using different substrates under standard assay conditions. Among various substrates, the purified PrASNase showed the maximum activity in the presence of its native substrate, L-asparagine, whereas little enzyme activity was observed when the D-asparagine was used as a substrate (Fig. 4A). Furthermore, PrASNase did not show any activity towards L-glutamine and other substrates. No activity of PrASNase towards glutamine indicated the enzyme to be glutaminase free, one major desirable feature. The kinetic parameters of PrASNase with L-asparagine as substrate, determined using the Lineweaver-Burk plot, showed the K_m_ and V_max_ of the purified PrASNase were 9.49×10^-3^ M and 25.13 IU mL^-1^ min^-1^ (10285 nanomoles mL^-1^ min^-1^ in terms of product formation), respectively.

**Fig. 4.**
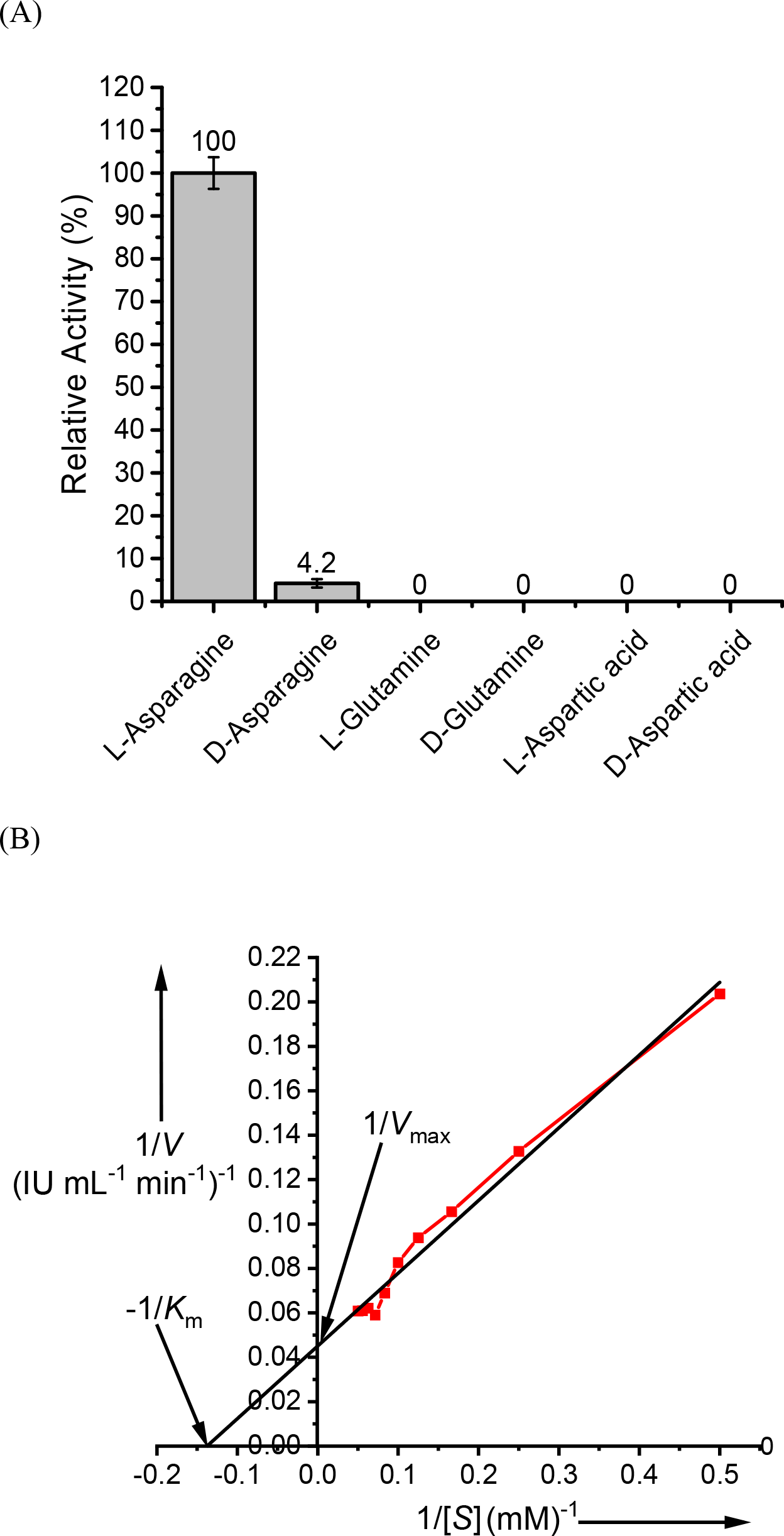
Substrate specificity and kinetic parameters of the purified PrASNase (A) Enzyme activity in the presence of different substrates, and (B) Lineweaver-Burk plot for the determination of kinetic parameters of the enzyme.

The turnover number (k_cat_) of the enzyme, defined as the formation of a product from the substrate by one site of enzyme per second, was calculated as 3.01×10^3^ s^-1^ (Fig. 4B). As the K_m_ of the enzyme reflected the affinity towards its substrate, the lower K_m_ value of the enzyme indicated a robust binding affinity of the L-asparaginase towards the substrate L-asparagine. The V_max_ indicated the reaction rate under which the enzyme activity was exhausted or saturated with substrate. The K_m_ value of PrASNase reported in the present study was similar to the K_m_ value of L-asparaginase produced from the Streptomyces fradiae NEAE-82 [36]. The reported K_m_ values of L-asparaginase produced from Cladosporium sp., Bacillus licheniformis RAM-8, and Pectobacterium carotovorum MTCC 1428 were lower than the K_m_ value of PrASNase [3,37,38].

### 3.5. Circular dichroism (CD) spectroscopy

The far-UV CD spectra of purified PrASNase monitored between 200-260 nm wavelengths showed negative ellipticities between the wavelengths 205-240 nm (Fig. 5A). The data obtained from this far-UV CD was analyzed on the BestSel server (http://bestsel.elte.hu/index.php) [39], which showed that the purified PrASNase in its native form consisted of 30.9% α-helix and 69.1% other structural content. The pattern of far UV spectra of purified PrASNase revealed its similarity with far-UV CD spectra of L-asparaginase produced from Bacillus licheniformis RAM-8 and E. coli type II asparaginase [3,40,41].

**Fig. 5.**
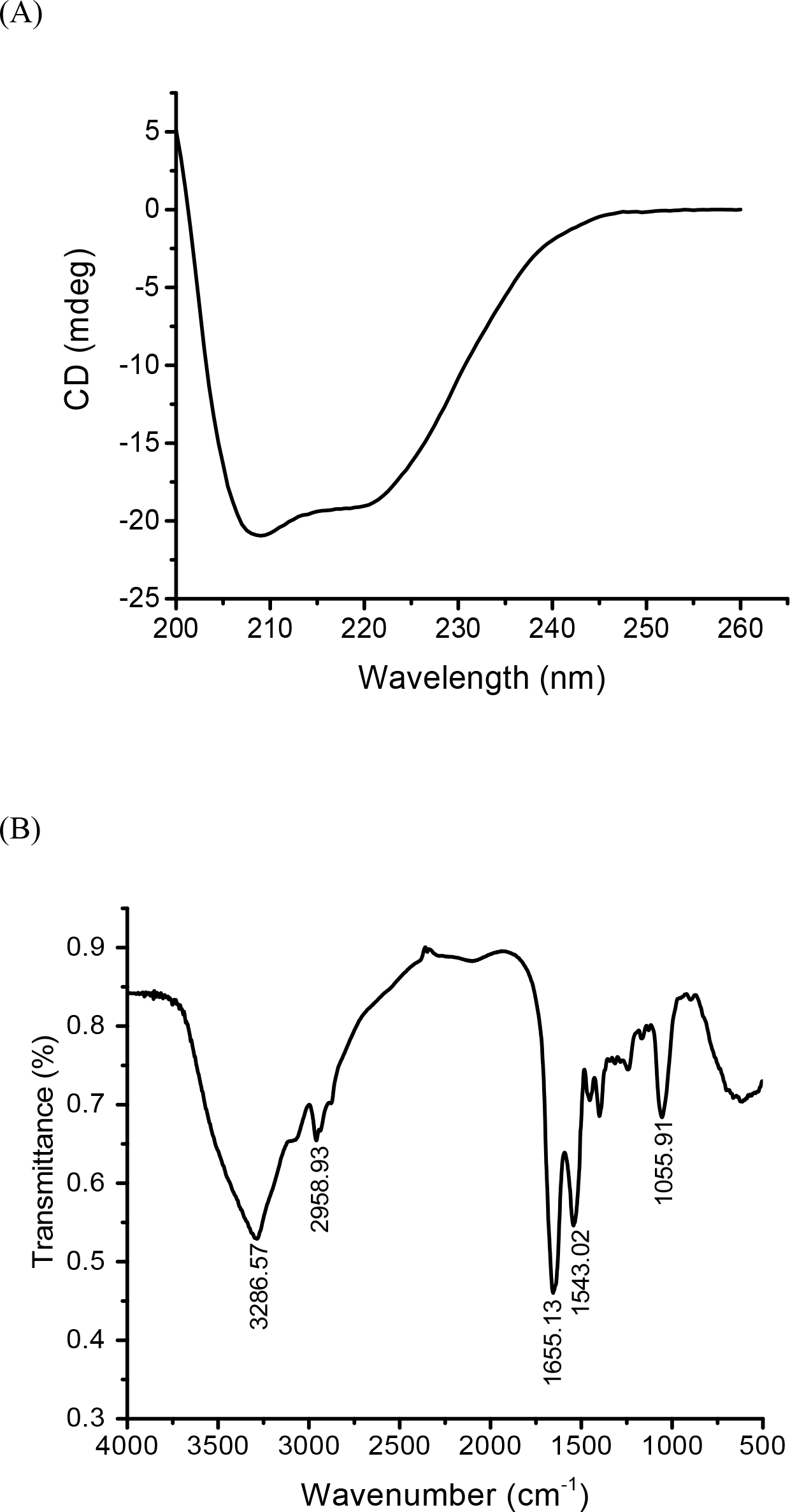
(A) The far-UV spectra of the purified PrASNase and (B) The FTIR spectrum of the purified PrASNase.

### 3.6. Fourier transform infrared spectroscopy (FTIR)

The presence of chemical and functional groups in the purified PrASNase was investigated using FTIR analysis. The spectrum obtained after FTIR revealed a peak at 3286.57 cm^-1,^ indicating the presence of N-H and O-H bonds. The stretching of the peak from 3286.57 cm^-1^ to 2958.93 cm^-1^ indicated the presence of –NH^3+^ interactions. The peaks at 1655.13 cm^-1^ and 1543.02 cm^-1^ in the spectrum reflected the presence of amide I and amide II bands, respectively, in the purified PrASNase. The stretching of the peak at 1055.91 cm^-1^ showed P = O groups in the PrASNase (Fig. 5B). Furthermore, the current FTIR spectrum of the purified PrASNase resembled the FTIR spectrum of two previously reported L-asparaginases [42, 43].

### 3.7. Oligomeric state analysis using SEC

The glutaraldehyde-treated PrASNase and untreated PrASNase were stored at 4°C, and SEC profiles of both samples were compared at 0 h, 24 h, and 48 h. The glutaraldehyde-treated sample showed a prominent peak at the volumes of 14.5 to 14.6 mL from 0 h to 48 h, which corresponds to the ∼72 kDa, which was the dimeric molecular weight of PrASNase (Fig. 6A, 6C, and 6E). On the other hand, the untreated PrASNase showed a natively dimeric peak (i.e., at a volume of 14.6 mL) at 0 h, both dimeric and monomeric peaks (i.e., at the volume of 14.7 and 16.28 mL) at 24 h, and prominent monomeric peak (i.e., at a volume of 16.30 mL) at 48 h (Fig. 6B, 6D, and 6F). This study showed the PrASNase was natively dimeric and dissociated in monomers upon storage at 4°C.

**Fig. 6.**
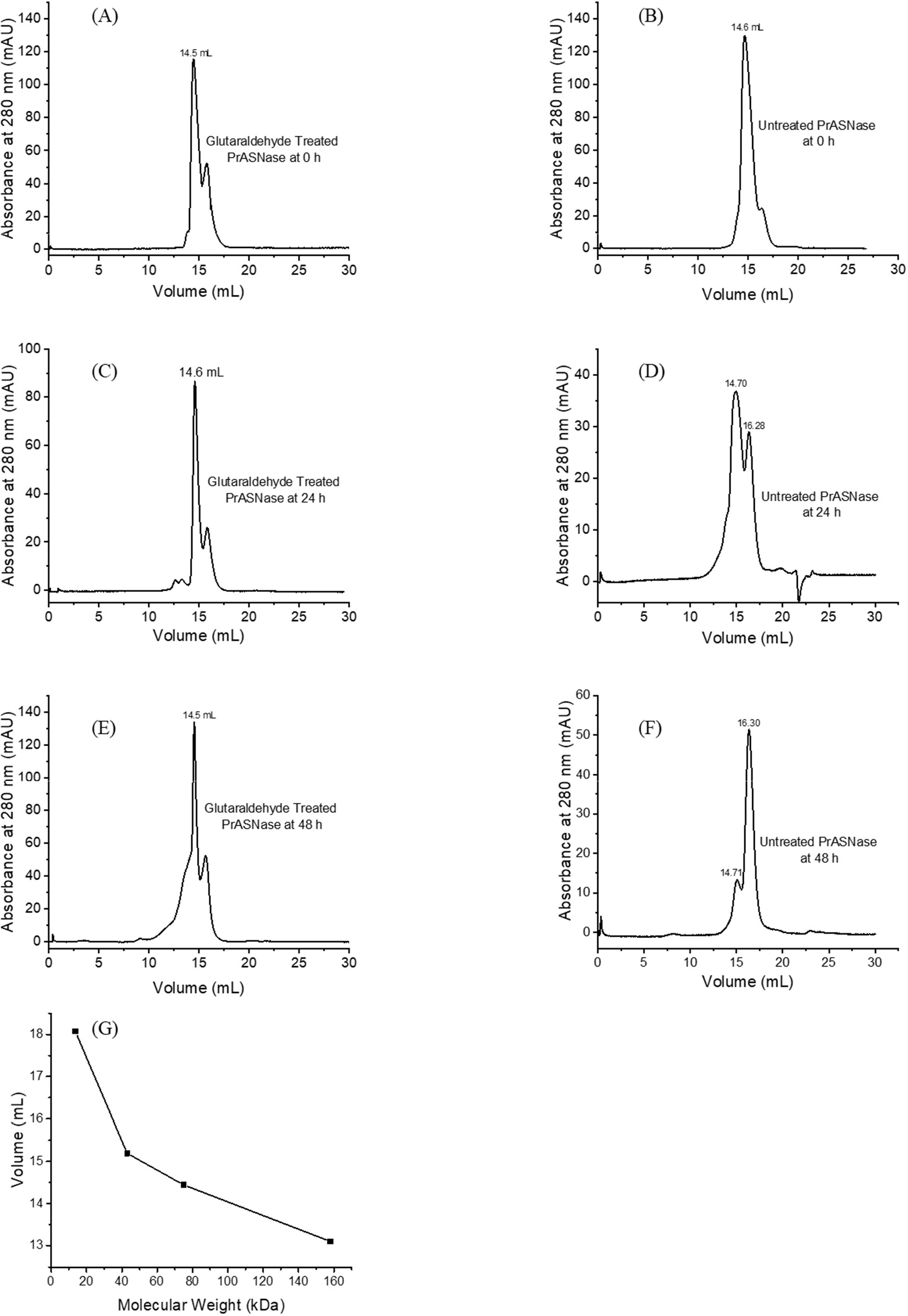
SEC profiles of PrASNase treated with glutaraldehyde and untreated at different time intervals (A) Glutaraldehyde treated PrASNase at 0 h, (B) Untreated PrASNase at 0 h, (C) Glutaraldehyde treated PrASNase at 24 h, (D) Untreated PrASNase at 24 h, (E) Glutaraldehyde treated PrASNase at 48 h, (F) Untreated PrASNase at 48 h, and (G) Molecular weight standards profile.

### 3.8. Small Angle X-ray Scattering (SAXS)

Previous investigations using crystallography, SAXS, and molecular modeling in the case of L- asparaginase from Pyrococcus furiosus have shown the protein (L-asparaginase) to be mainly dimer and holding its dimeric structure despite the lack of a large interconnecting linker [44]. SAXS datasets were acquired for PrASNase, which showed a monomeric profile in SEC analysis at a concentration close to 2 mg mL-1, shown in Fig. 7, at a range of temperatures. The downward trend of the data as q approaches 0 nm^-1^ indicates the presence of inter particulate effect in the sample, making the molecules repel each other (Fig. 7A). The behaviour was retained in protein molecules even at elevated temperatures. Normalized Kratky plots shown in Fig. 7B showed peak-like behaviour with maxima around 1.73 supporting the retention of globular scattering profile in the PrASNase molecules throughout the temperature range. The variation in R_g_ was plotted as a function of temperature in Fig. 7C. The trend indicated that PrASNase molecules underwent a rapid increase in size due to heat-induced association with the mid-point of transition around 37°C (310 K). This was supported by a systematic increase in the calculated I_0_ values for the datasets (Supplementary Table S2). Using the reference value at 10°C (283 K), I_0_ values estimated the molecules’ association status. The PrASNase was mostly in monomeric form till 30°C (303 K), and with a rise in temperature, the formation of some intermediates (mixed order) started at 37°C (310 K). Interestingly, unlike those seen for thermophilic L-asparaginase, the reduction in experimental temperature did not induce a decrement of the scattering particles in the solution, supporting the irreversibility of the heat-induced association of the PrASNase molecules [45]. To explore if the heat-induced association can be affected by osmolytes, 300 mM arginine and 2% sucrose were added to the protein (and buffer) by dialysis, and SAXS experiments were repeated. The mapping of R_g_ values showed that both osmolytes delay the onset of association of the PrASNase molecules but do not play a role in inducing reversibility to a lower-order state with a lowering of temperature. This information will be borne in mind during setting up future experiments like attempts to crystallize this protein.

**Fig. 7.**
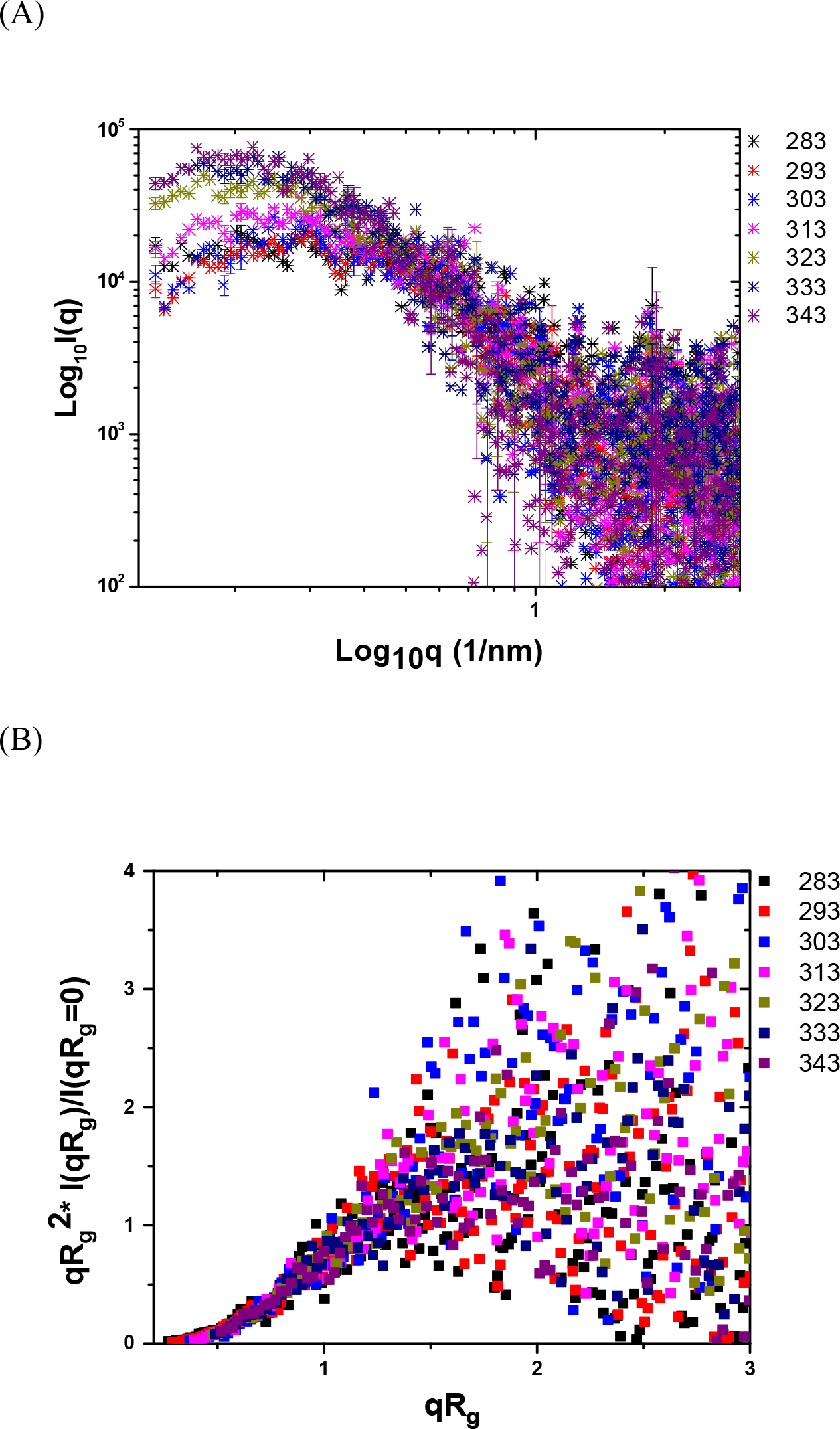
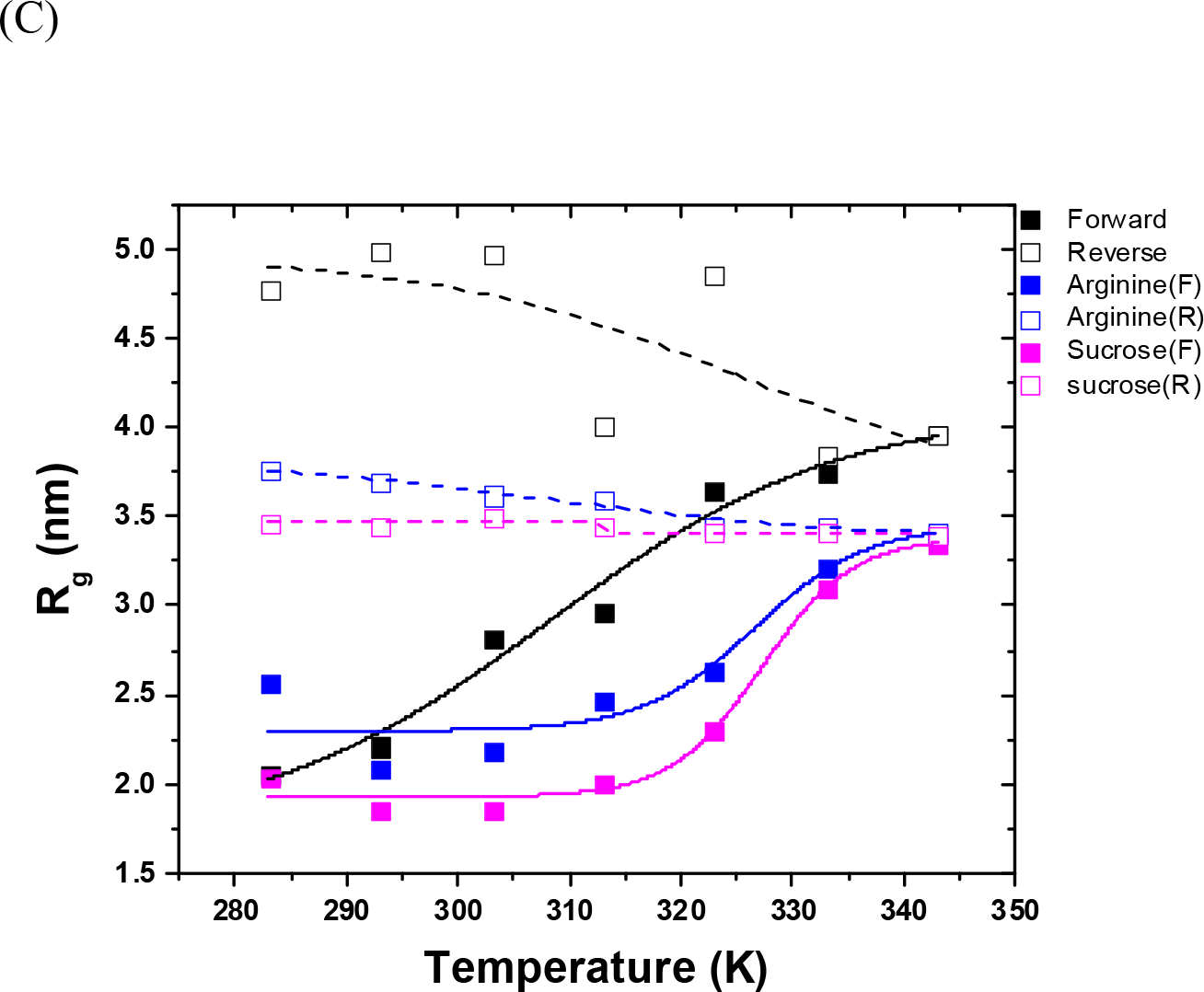
SAXS datasets for PrASNase (A) The SAXS I(q) profiles of PrASNase at different temperatures (K), (B) Normalized Kratky plot of the datasets, and (C) The variation in the radius of gyration (R_g_) of apo-protein and on the addition of arginine and sucrose, as a function of forward and reversal of temperature.

As mentioned in the methods, shape restoration of the PrASNase molecules was done using chain-ensemble modeling protocols using online programs of ATSAS and averaged using the offline DAMAVER program. The same protocol was used for generating shapes of dimer and tetramer of PrASNase protein. In Fig. 8A, the computed normalized spatial disposition (NSD) values are listed below each model as a reflection of model similarity. Alternative rotated views of the same are shown in Supplementary Fig. S1. A value close to 1 indicates the identical nature of the models being compared. The dummy residue shape was superimposed with the model generated, considering one chain from PDB ID 3NTX as a template by aligning their inertial axes. χ^2^ value was computed between the theoretical SAXS profile of the final dummy residue model and the model of PrASNase generated by the Swiss modeler server, and later for dimer or tetramer using symmetry mates of the crystal structure. Again, a value of 1 is supportive of a similar nature. The models and superimpositions shown in Fig. 8A indicated that the PrASNase protein adopts a monomeric state at lower temperatures in solution, which forms an antiparallel dimer at increased temperatures. This dimer then associates to form a tetramer. The PrASNase was catalytically active against L-asparagine when present in a monomeric state but did not show any activity when adopting a higher oligomeric state upon the increase in temperature (data not shown). The repetitive manner of association is summarized in the schematic in Fig. 8B. It also indicates that the active site of the protein may get occluded with an increase in temperature, and thus, in states or temperatures forming higher-order association than dimer, the total activity of the protein may drastically reduce. This “side-ways” association profile is in contrast to the temperature-induced end-to-end association of L-asparaginase from P. furiosus (PfLAsp) when heated [46]. The differential relevance of this contrasting behaviour to the shape-function of this enzyme is that while PrASNase displayed diminished activity with heating, while thermophile PfLAsp showed enhanced activity at elevated temperatures than ambient or physiological ones.

**Fig. 8.**
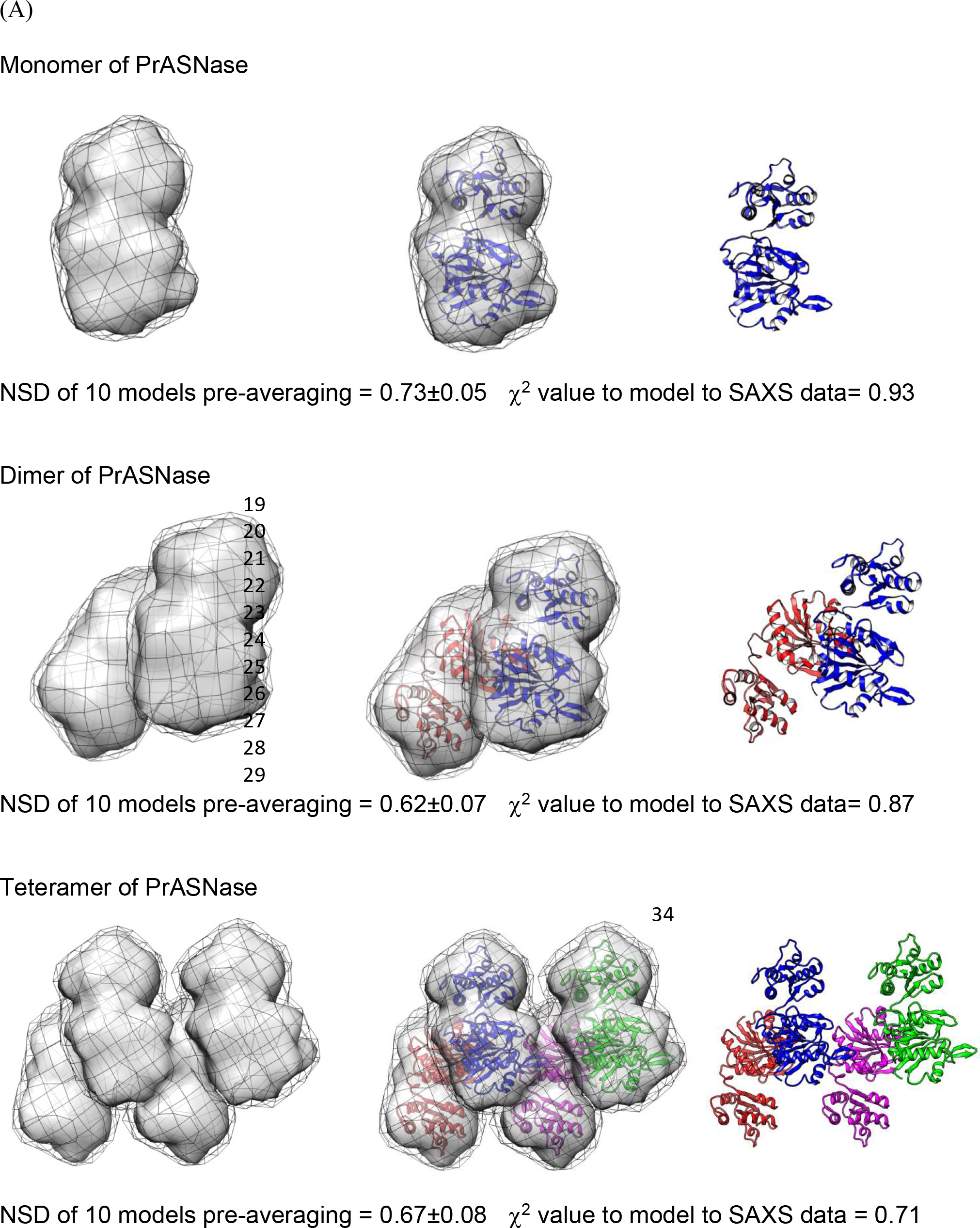
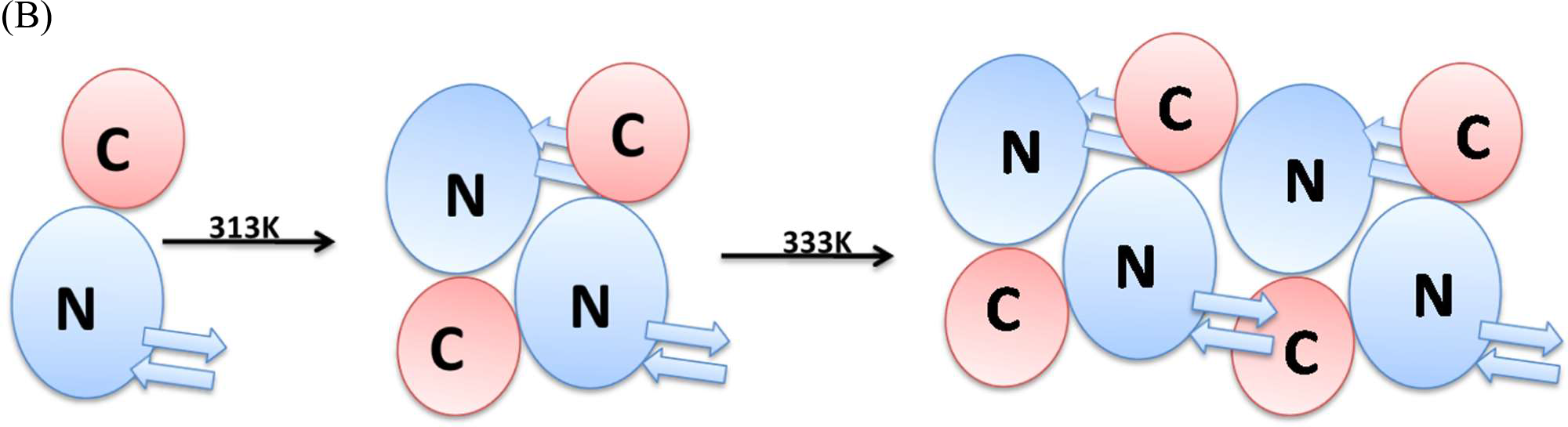
SAXS data-based scattering shape models (A) Monomer, dimer, and tetramer form of PrASNase (molecular maps), and (B) Schematic of anti-parallel association of PrASNase molecules with heating.

### 3.9. Sedimentation velocity analytical ultracentrifugation (SV-AUC)

SV-AUC is an orthogonal method to SEC where samples are not subjected to extraneous conditions like solid or mobile phases. This analysis facilitated the accurate determination of the molecular mass and oligomeric state of proteins present in a solution based on its sedimentation coefficient value [47]. The SV-AUC analysis (carried out at 25°C-30°C) of the purified catalytically active PrASNase samples (stored at 4°C) revealed that 94.9% of PrASNase was present as a monomer with a molecular weight of 35.6 kDa and sedimentation coefficient value of 2.8 while only 4.8% population of protein was present as a dimer with a molecular weight of 71.1 kDa and sedimentation coefficient value of 5.7 (Fig. 9). It could be concluded from this SV-AUC analysis of PrASNase that most protein species were monomer in solution, which was natively dimeric and catalytically active as the protein from the same sample showed high affinity towards its substrate in enzyme kinetics studies. Although the PrASNase was possibly dissociated into monomers after a few days of storage at 4°C, its monomeric form and catalytic activities were maintained at a temperature of around 37°C.

**Fig. 9.**
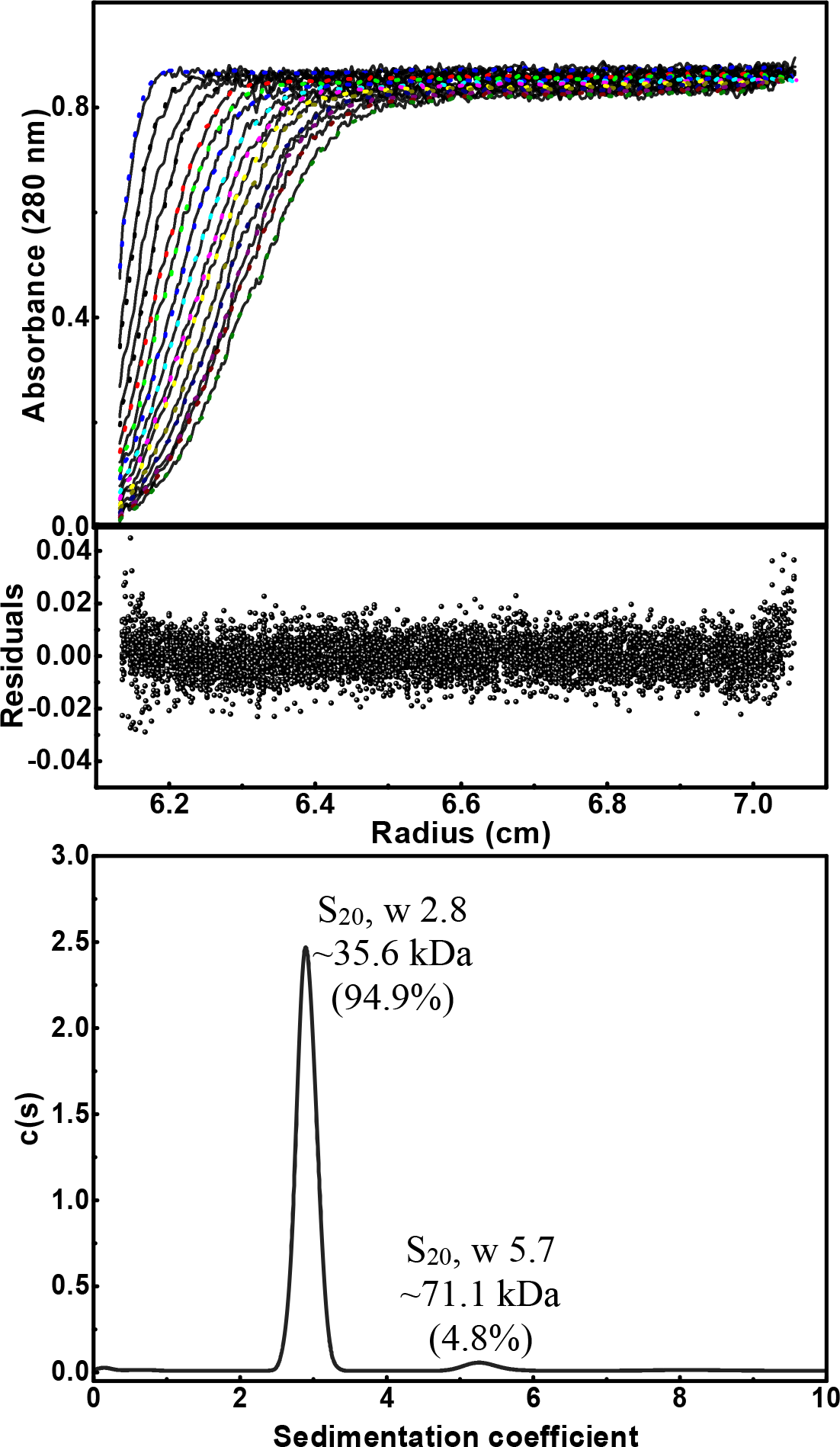
Sedimentation velocity (SV) for determining the oligomeric state of the purified PrASNase by using SV-AUC.

### 3.10. Anti-cancerous effect of PrASNase

The anti-cancerous effect of the monomeric PrASNase was investigated against the WIL2-S and TF-1.28 cancer cell lines by using Alamar blue dye in the cell-based inhibition method. The profiles of proliferation inhibition profiles of cancer cell lines, in terms of RFU, were generated after 24 h of treatment with PrASNase (Fig. 10). The IC_50_ value of 7.4 µg mL^-1^ and 5.6 µg mL^-1^ of the monomeric PrASNase was highly effective in retarding or inhibiting the growth of WIL2- S and TF-1.28 cancer cell lines, respectively. The L-asparaginase produced from Bacillus licheniformis has been reported to have a cytotoxic effect towards HepG-2, MCF-7, and HCT- 116 cells with IC_50_ values of 11.66 µg mL^-1^, 14.55 µg mL^-1^, and 17.02 µg mL^-1^, respectively [48]. Another asparaginase from Bacillus licheniformis RAM-8 with IC_50_ values of 0.22 IU, 0.15 IU, and 0.78 IU was greatly effective against the cancerous cells E6-1, K-562, and MCF-7, respectively [3]. The asparaginase from wild-type Erwinia with IC_50_ values of 0.33 IU mL^-1^ and 0.16 IU mL^-1^ was reported to be effective against the LOUCY and SUP-B15, respectively, and that from wild-type E. coli asparaginase with IC_50_ values of 0.35 IU mL^-1^ and 0.22 IU mL^-1^ were effective against similar cell lines [49]. The L-asparaginase produced from Zymomonas mobilis and Pyrococcus furiosus also showed significant anti-cancerous activity against various cell lines, including Reh cells, HL60 cells, K-562 cells, and MCF-7 breast carcinoma cell lines [14, 22]. Another L-asparaginase from Fusarium equiseti AHMF4 showed an anti-cancerous activity against cervical epitheloid carcinoma (2.0 µg mL^-1^), colorectal carcinoma (HCT-116) (8.26 µg mL^-1^), breast adenocarcinoma (MCF-7) (22.8 µg mL^-1^), epidermoid larynx carcinoma (Hep-2) (5.0 µg mL^-1^), and hepatocellular carcinoma (HepG-2) (12.40 µg mL^-1^) [50].

**Fig. 10.**
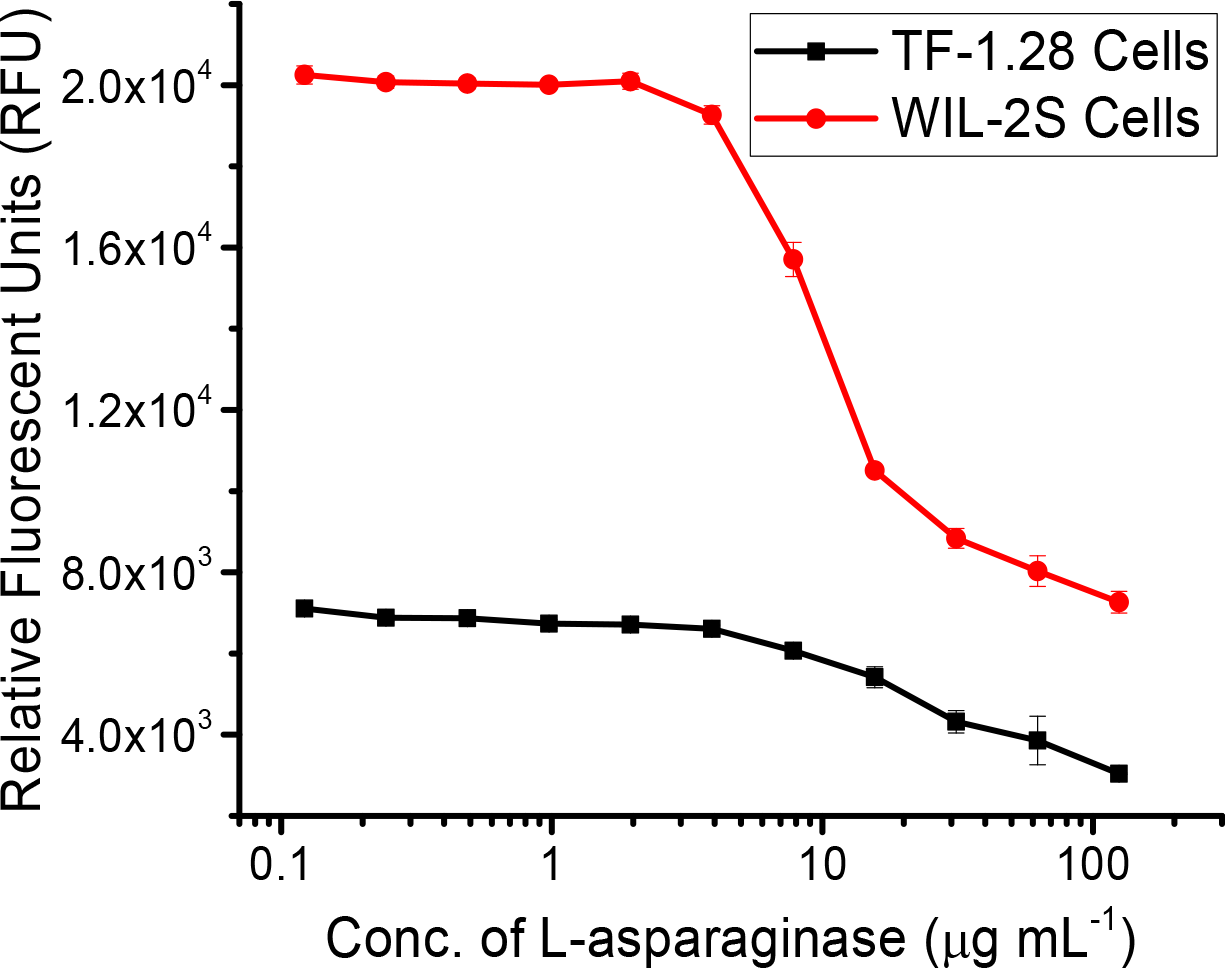
Anti-cancerous effect of the purified PrASNase on the WIL2-S and TF-1.28 leukemic cell lines.

## 4. Conclusion

In the current study, the alignment study showed the L-asparaginase, PrASNase, to consist of the conserved motifs upon alignment with EcASNase, EwASNase, BsASNase, and PASNase. The molecular docking speculated that the asparagine binds to serine, alanine, and glutamine in the binding pocket. The PrASNase was highly specific for its substrate L-asparagine with ideal values of enzyme kinetic parameters and was found to contain α-helix and other structures. The FTIR analysis showed the presence of N-H and O-H bonds, –NH^3+^ interactions, amide I and II bands, and P = O groups in the purified enzyme. The L-asparaginase (PrASNase) stored in 20 mM Tris-Cl buffer (200 mM NaCl and pH 8.0) at 4°C for 48 h was found to mainly consist of monomeric species, whereas it oligomerized to the higher state as the temperature increased. SAXS analysis showed the protein to be mostly monomeric till 37°C and catalytically active at that temperature under enzyme assay conditions. As discussed above, the side-ways association of this enzyme upon heating is in sharp contrast with the end-to-end spiral order of association for thermophilic P. furiosus L-asparaginase [46]. In the latter, the dimeric association is the building block of the higher-order association that disassembles upon heating withdrawal. At the same time, for PrASNase, the heat-induced association is not reversible. Even the addition of osmolytes did not alter this behaviour. Possibly the higher-order packing is different between the two enzymes in the form that the active sites of the enzymes are differentially accessible. The higher-order heat-induced packing in PrASNase possibly renders its active site inactive, which reflects the decrement of its activity with heating. The finding of functional L-asparaginase in monomeric form is considered highly significant and novelty of PrASNase as it is first reported with experimental evidence. Further studies, such as protein modeling and complete structure elucidation, will help decipher the factors responsible for L-asparaginase activity in monomeric form. The purified monomeric PrASNase showed a significant anti-cancerous effect against leukemic cell lines WIL2-S and TF-1.28. The physicochemical characterization and anti-cancerous study of the novel L-asparaginase (PrASNase) showed its potential as a candidate for pharmaceutical and other industrial applications.

## Acknowledgement

The authors Kanti Nandan Mihooliya and Jitender Nandal are thankful to the Council of Scientific and Industrial (CSIR) and the Department of Biotechnology (DBT). Special thanks to Dr Alka Kumari for helping with alignment studies.

## Author contributions

Kanti N. Mihooliya: Conceptualization, Visualization, Investigation, Writing – Original Manuscript.

Jitender Nandal: Visualization and Investigation. Nidhi Kalidas: Investigation of SAXS experiments.

Ashish: Conceptualization, Investigation, and Writing of SAXS experiments in the Manuscript.

Subhash Chand: Investigation of Biological assay in the Manuscript. Mani S. Bhattacharyya: Manuscript Review.

Debendra K. Sahoo: Conceptualization, Investigation, Visualization, Supervision, Writing - Review and Editing of the Manuscript.

## References

1. R. v. Mahajan, S. Saran, K. Kameswaran, V. Kumar, R.K. Saxena, Efficient production of L-asparaginase from Bacillus licheniformis with low-glutaminase activity: optimization, scale up and acrylamide degradation studies, Bioresour Technol. 125 (2012) 11–16. https://doi.org/10.1016/j.biortech.2012.08.086.

2. R. v. Mahajan, K.N. Mihooliya, S. Saran, R.K. Saxena, L-Asparaginase from Bacillus sp. RKS-20: Process optimization and application in the inhibition of acrylamide formation in fried foods, J Proteins Proteom. 5 (2014) 125–132.

3. R. v. Mahajan, V. Kumar, V. Rajendran, S. Saran, P.C. Ghosh, R.K. Saxena, Purification and characterization of a novel and robust L-asparaginase having low-glutaminase activity from Bacillus licheniformis: In vitro evaluation of anti-cancerous properties, PLoS One. 9 (2014). https://doi.org/10.1371/journal.pone.0099037.

4. J. Lubkowski, G.J. Palm, G.L. Gilliland, C. Derst, K.H. Röhm, A. Wlodawer, Crystal structure and amino acid sequence of Wolinella succinogenes L-asparaginase, Eur J Biochem. 241 (1996) 201–207. http://www.ncbi.nlm.nih.gov/pubmed/8898907.

5. J. Lubkowski, A. Wlodawer, Structural and biochemical properties of L-asparaginase, FEBS J. 288 (2021) 4183–4209. https://doi.org/10.1111/FEBS.16042.

6. K.B. McCredie, D.H. Ho, E.J. Freireich, L-Asparaginase for the treatment of cancer, CA Cancer J Clin. 23 (1973) 220–227.

7. T. Ohnuma, J.F. Holland, A. Freeman, L.F. Sinks, Biochemical and pharmacological studies with asparaginase in man, Cancer Res. 30 (1970) 2297–2305.

8. M. Chiu, G. Taurino, M.G. Bianchi, M.S. Kilberg, O. Bussolati, Asparagine synthetase in cancer: beyond acute lymphoblastic leukemia, Front Oncol. 9 (2020). https://doi.org/10.3389/fonc.2019.01480.

9. Z. Ciesarová, E. Kiss, P. Boegl, Impact of L-asparaginase on acrylamide content in potato products, Journal of Food and Nutrition Research. 45 (2006) 141–146. https://doi.org/10.1016/B978-0-12-802832-2.00021-8.

10. F. Xu, M.J. Oruna-Concha, J.S. Elmore, The use of asparaginase to reduce acrylamide levels in cooked food, Food Chem. 210 (2016) 163–171. https://doi.org/10.1016/j.foodchem.2016.04.105.

11. M. Duval, S. Suciu, A. Ferster, X. Rialland, B. Nelken, P. Lutz, Y. Benoit, A. Robert, A.M. Manel, E. Vilmer, J. Otten, N. Philippe, Comparison of Escherichia coli- asparaginase with Erwinia-asparaginase in the treatment of childhood lymphoid malignancies: Results of a randomized European Organisation for Research and Treatment of Cancer - Children’s leukemia group phase 3 trial, Blood. 99 (2002) 2734– 2739. https://doi.org/10.1182/blood.V99.8.2734.

12. L.P. Brumano, F.V.S. da Silva, T.A. Costa-Silva, A.C. Apolinário, J.H.P.M. Santos, E.K. Kleingesinds, G. Monteiro, C. de O. Rangel-Yagui, B. Benyahia, A.P. Junior, Development of L-asparaginase biobetters: Current research status and review of the desirable quality profiles, Front Bioeng Biotechnol. 6 (2019). https://doi.org/10.3389/fbioe.2018.00212.

13. J.C.F. Nunes, R.O. Cristóvão, M.G. Freire, V.C. Santos-Ebinuma, J.L. Faria, C.G. Silva, A.P.M. Tavares, Recent strategies and applications for L-asparaginase confinement, Molecules. 25 (2020). https://doi.org/10.3390/molecules25245827.

14. S. Bansal, A. Srivastava, G. Mukherjee, R. Pandey, A.K. Verma, P. Mishra, B. Kundu, Hyperthermophilic asparaginase mutants with enhanced substrate affinity and antineoplastic activity: structural insights on their mechanism of action, The FASEB Journal. 26 (2012) 1161–1171. https://doi.org/10.1096/fj.11-191254.

15. A.M. Schalk, H.A. Nguyen, C. Rigouin, A. Lavie, Identification and structural analysis of an L-asparaginase enzyme from guinea pig with putative tumor cell killing properties, Journal of Biological Chemistry. 289 (2014) 33175–33186. https://doi.org/10.1074/jbc.M114.609552.

16. Z.-J. Dai, Y.-Q. Huang, Y. Lu, Efficacy and safety of PEG-asparaginase versus E. coli L-asparaginase in Chinese children with acute lymphoblastic leukemia: a meta-analysis, Transl Pediatr. 10 (2021) 244–255. https://doi.org/10.21037/tp-20-178.

17. S. Bansal, D. Gnaneswari, P. Mishra, B. Kundu, Structural stability and functional analysis of L-asparaginase from Pyrococcus furiosus, Biochemistry (Moscow). 75 (2010) 375–381. https://doi.org/10.1134/S0006297910030144.

18. D.M. Charbonneau, A. Aubé, N.M. Rachel, V. Guerrero, K. Delorme, J. Breault-Turcot, J.F. Masson, J.N. Pelletier, Development of Escherichia coli asparaginase II for immunosensing: A trade-off between receptor density and sensing efficiency, ACS Omega. 2 (2017) 2114–2125. https://doi.org/10.1021/acsomega.7b00110.

19. D.K. Garg, B. Kundu, Hyperthermophilic L-asparaginase bypasses monomeric intermediates during folding to retain cooperativity and avoid amyloid assembly, Arch Biochem Biophys. 622 (2017) 36–46. https://doi.org/10.1016/j.abb.2017.04.010.

20. A.A. Pritsa, D.A. Kyriakidis, L-Asparaginase of Thermus thermophilus: Purification, properties and identification of essential amino acids for its catalytic activity, Mol Cell Biochem. 216 (2001) 93–101. https://doi.org/10.1023/A:1011066129771.

21. G.A. Kotzia, N.E. Labrou, Cloning, expression and characterisation of Erwinia carotovora L-asparaginase, J Biotechnol. 119 (2005) 309–323. https://doi.org/10.1016/j.jbiotec.2005.04.016.

22. K. Einsfeldt, I.C. Baptista, J.C.C.V. Pereira, I.C. Costa-Amaral, E.S. Da Costa, M.C.M. Ribeiro, M.G.P. Land, T.L.M. Alves, A.L. Larentis, R.V. Almeida, Recombinant L-asparaginase from Zymomonas mobilis: A potential new antileukemic agent produced in Escherichia coli, PLoS One. 11 (2016). https://doi.org/10.1371/journal.pone.0156692.

23. K.N. Mihooliya, J. Nandal, A. Kumari, S. Nanda, H. Verma, D.K. Sahoo, Studies on efficient production of a novel L-asparaginase by a newly isolated Pseudomonas resinovorans IGS-131 and its heterologous expression in Escherichia coli, 3 Biotech. 10 (2020). https://doi.org/10.1007/s13205-020-2135-4.

24. A. Imada, S. Igarasi, K. Nakahama, M. Isono, Asparaginase and glutaminase activities of micro-organisms, J Gen Microbiol. 76 (1973) 85–99. https://doi.org/10.1099/00221287-76-1-85.

25. K.N. Mihooliya, J. Nandal, L. Swami, H. Verma, L. Chopra, D.K. Sahoo, A new pH indicator dye-based method for rapid and efficient screening of L-asparaginase producing microorganisms., Enzyme Microb Technol. 107 (2017) 72–81. https://doi.org/10.1016/j.enzmictec.2017.08.004.

26. N. Chakravarty, N. Priyanka, J. Singh, R. Singh, A potential type-II L-asparaginase from marine isolate Bacillus australimaris NJB19: Statistical optimization, in silico analysis and structural modeling, Int J Biol Macromol. 174 (2021) 527–539. https://doi.org/10.1016/J.IJBIOMAC.2021.01.130.

27. H. Saeed, H. Ali, H. Soudan, A. Embaby, A. El-Sharkawy, A. Farag, A. Hussein, F. Ataya, Molecular cloning, structural modeling and production of recombinant Aspergillus terreus L-asparaginase in Escherichia coli, Int J Biol Macromol. 106 (2018) 1041–1051. https://doi.org/10.1016/j.ijbiomac.2017.08.110.

28. I.M. Costa, L. Schultz, B. de Araujo Bianchi Pedra, M.S.M. Leite, S.H.P. Farsky, M.A. de Oliveira, A. Pessoa, G. Monteiro, Recombinant L-asparaginase 1 from Saccharomyces cerevisiae: An allosteric enzyme with antineoplastic activity, Sci Rep. 6 (2016). https://doi.org/10.1038/srep36239.

29. W. Zheng, C. Zhang, Y. Li, R. Pearce, E.W. Bell, Y. Zhang, Folding non-homologous proteins by coupling deep-learning contact maps with I-TASSER assembly simulations, Cell Reports Methods. 1 (2021). https://doi.org/10.1016/J.CRMETH.2021.100014.

30. G.C.P. van Zundert, J.P.G.L.M. Rodrigues, M. Trellet, C. Schmitz, P.L. Kastritis, E. Karaca, A.S.J. Melquiond, M. van Dijk, S.J. de Vries, A.M.J.J. Bonvin, The HADDOCK2.2 web server: User-friendly integrative modeling of biomolecular complexes, J Mol Biol. 428 (2016) 720–725. https://doi.org/10.1016/J.JMB.2015.09.014.

31. H. Lineweaver, D. Burk, The determination of enzyme dissociation constants, J Am Chem Soc. 56 (1934) 658–666. https://doi.org/10.1021/ja01318a036.

32. M.D. Badmalia, P. Sharma, S.P.S. Yadav, S. Singh, N. Khatri, R. Garg, Ashish, Bonsai gelsolin survives heat induced denaturation by forming β-amyloids which leach out functional monomer, Sci Rep. 8 (2018). https://doi.org/10.1038/s41598-018-30951-3.

33. K. Dhiman, S.K. Nath, Ashish, Monomeric human soluble CD4 dimerizes at physiological temperature: VTSAXS data based modeling and screening of retardant molecules, J Biomol Struct Dyn. (2020). https://doi.org/10.1080/07391102.2020.1771422.

34. S. Agrawal, U.K. Jana, N. Kango, Heterologous expression and molecular modelling of L-asparaginase from Bacillus subtilis ETMC-2, Int J Biol Macromol. 192 (2021) 28–37. https://doi.org/10.1016/J.IJBIOMAC.2021.09.186.

35. M. Sanches, S. Krauchenco, I. Polikarpov, Structure, substrate complexation and reaction mechanism of bacterial Asparaginases, Curr Chem Biol. 1 (2012) 75–86. https://doi.org/10.2174/2212796810701010075.

36. N.E.A. El-Naggar, S.F. Deraz, H.M. Soliman, N.M. El-Deeb, S.M. El-Ewasy, Purification, characterization, cytotoxicity and anticancer activities of L-asparaginase, anti-colon cancer protein, from the newly isolated alkaliphilic Streptomyces fradiae NEAE-82, Sci Rep. 6 (2016). https://doi.org/10.1038/srep32926.

37. S. Kumar, V. Venkata Dasu, K. Pakshirajan, Purification and characterization of glutaminase-free L-asparaginase from Pectobacterium carotovorum MTCC 1428, Bioresour Technol. 102 (2011) 2077–2082. https://doi.org/10.1016/j.biortech.2010.07.114.

38. N.S. Mohan Kumar, H.K. Manonmani, Purification, characterization and kinetic properties of extracellular L-asparaginase produced by Cladosporium sp., World J Microbiol Biotechnol. 29 (2013). https://doi.org/10.1007/s11274-012-1213-0.

39. A. Micsonai, F. Wien, É. Bulyáki, J. Kun, É. Moussong, Y.H. Lee, Y. Goto, M. Réfrégiers, J. Kardos, BeStSel: a web server for accurate protein secondary structure prediction and fold recognition from the circular dichroism spectra, Nucleic Acids Res. 46 (2018) W315–W322. https://doi.org/10.1093/NAR/GKY497.

40. A.K. Upadhyay, A. Singh, K.J. Mukherjee, A.K. Panda, Refolding and purification of recombinant L-asparaginase from inclusion bodies of E. coli into active tetrameric protein, Front Microbiol. 5 (2014). https://doi.org/10.3389/fmicb.2014.00486.

41. A. P. Singhvi, J. Verma, N. Panwar, T.Q. Wani, A. Singh, M. Qudratullah, A. Chakraborty, A. Saneja, D.P. Sarkar, A.K. Panda, Molecular attributes associated with refolding of inclusion body proteins using the freeze-thaw method, Front Microbiol. 12 (2021). https://doi.org/10.3389/fmicb.2021.618559.

42. B. Meena, L. Anburajan, N.V. Vinithkumar, D. Shridhar, R.V. Raghavan, G. Dharani, R. Kirubagaran, Molecular expression of L-asparaginase gene from Nocardiopsis alba NIOT-VKMA08 in Escherichia coli: A prospective recombinant enzyme for leukaemia chemotherapy, Gene. 590 (2016) 220–226. https://doi.org/10.1016/J.GENE.2016.05.003.

43. E. Bahreini, K. Aghaiypour, R. Abbasalipourkabir, A.R. Mokarram, M.T. Goodarzi, M. Saidijam, Preparation and nanoencapsulation of L-asparaginase II in chitosan- tripolyphosphate nanoparticles and in vitro release study, Nanoscale Res Lett. 9 (2014) 1– 13. https://doi.org/10.1186/1556-276X-9-340.

44. R. Tomar, P. Sharma, A. Srivastava, S. Bansal, Ashish, B. Kundu, Structural and functional insights into an archaeal L-asparaginase obtained through the linker-less assembly of constituent domains, Acta Crystallogr D Biol Crystallogr. 70 (2014) 3187– 3197. https://doi.org/10.1107/S1399004714023414.

45. P. Sharma, N. Verma, P.K. Singh, S. Korpole, Ashish, Characterization of heat induced spherulites of lysozyme reveals new insight on amyloid initiation, Sci Rep. 6 (2016). https://doi.org/10.1038/srep22475.

46. P. Sharma, R. Tomar, S.S. Yadav, M.D. Badmalia, S.K. Nath, Ashish, B. Kundu, Heat induces end to end repetitive association in P. furiosus L-asparaginase which enables its thermophilic property, Sci Rep. 10 (2020). https://doi.org/10.1038/S41598-020-78877-Z.

47. Y.R. Gokarn, M. McLean, T.M. Laue, Effect of PEGylation on protein hydrodynamics, Mol Pharm. 9 (2012) 762–773. https://doi.org/10.1021/mp200470c.

48. S.A. Alrumman, Y.S. Mostafa, K.A. Al-izran, M.Y. Alfaifi, T.H. Taha, S.E. Elbehairi, Production and anticancer activity of an L-asparaginase from Bacillus licheniformis isolated from the red sea, Saudi Arabia, Sci Rep. 9 (2019). https://doi.org/10.1038/s41598-019-40512-x

49. H.A. Nguyen, Y. Su, J.Y. Zhang, A. Antanasijevic, M. Caffrey, A.M. Schalk, L. Liu, D. Rondelli, A. Oh, D.L. Mahmud, M.C. Bosland, A. Kajdacsy-Balla, S. Peirs, T. Lammens, V. Mondelaers, B. De Moerloose, S. Goossens, M.J. Schlicht, K.K. Kabirov, A. V. Lyubimov, B.J. Merrill, Y. Saunthararajah, P. Van Vlierberghe, A. Lavie, A novel L-asparaginase with low L-glutaminase coactivity is highly efficacious against both T- and B-cell acute lymphoblastic Leukemias In Vivo, Cancer Res. 78 (2018) 1549–1560. https://doi.org/10.1158/0008-5472.CAN-17-2106.

50. M.M.A.A. El-Gendy, M.F. Awad, F.S. El-Shenawy, A.M.A. El-Bondkly, Production, purification, characterization, antioxidant and antiproliferative activities of extracellular L-asparaginase produced by Fusarium equiseti AHMF4, Saudi J Biol Sci. 28 (2021) 2540– 2548. https://doi.org/10.1016/j.sjbs.2021.01.058.

