## Supplementary Material for "Assessment of structural behaviour of new L-asparaginase and SAXS data-based evidence for catalytic activity in its monomeric form"

Research Article

#Acedemy of Scientific and Innovative Research

*Corresponding author:

Dr. Debendra K. Sahoo

Biochemical Engineering Research and Process Development Centre,

CSIR-Institute of Microbial Technology

Chandigarh-160 036, India

**Supplementary Fig. S1**

Alternated views of SAXS data-based scattering shape models of Monomer, dimer, and tetramer form of PrASNase (molecular maps).

Monomer of PrASNase

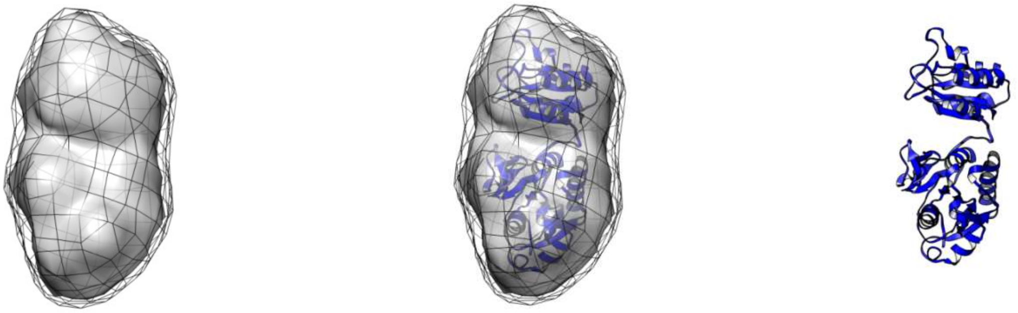

Dimer of PrASNase

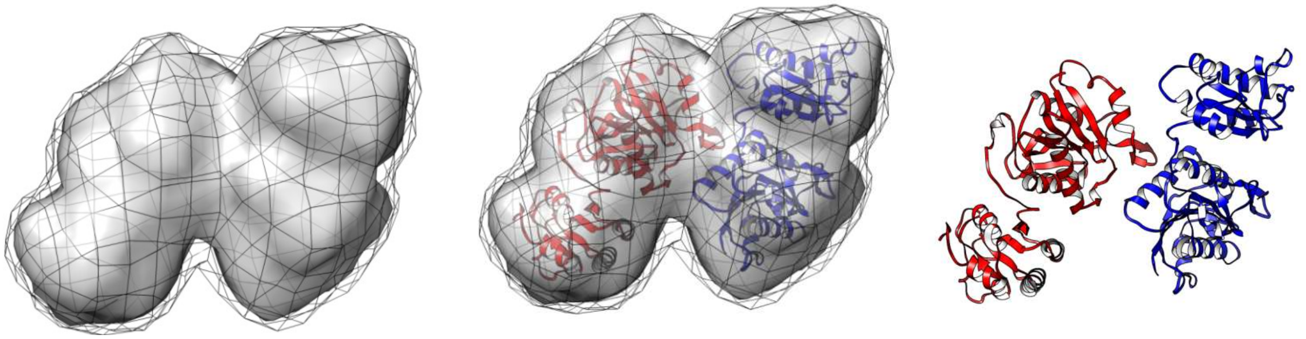

Tetramer of PrASNase

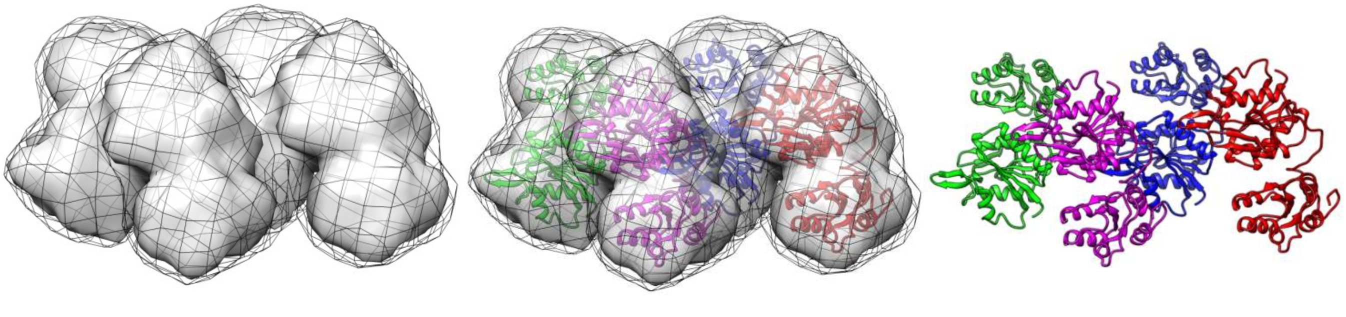

**Supplementary Table S1**

Different parameters deduced from the analysis of the SAXS datasets have been tabulated here for the Apo L-asparaginase protein and in the presence of 300 mM Arginine or 2% sucrose.

| Temperature (K) | Porod Exponent | $R_g$ (nm) | Intensity ($I_0$) | Association State* | Scattering Shape from Guinier plot (Association state) |
|---|---|---|---|---|---|
| Asparaginase | | | | | |
| 283 | 2.5 | 2.05 | 18036 | 1 | Globular (Monomer) |
| 293 | 2.6 | 2.21 | 18911 | 1.04 | Globular (Monomer) |
| 303 | 2.5 | 2.82 | 24564 | 1.3 | Globular (*Mixed order*) |
| 313 | 2.6 | 2.97 | 32388 | 1.7 | Globular (Mixed order) |
| 323 | 3.5 | 3.65 | 50385 | 2.8 | Globular (*Mixed order*) |
| 333 | 3.7 | 3.74 | 68019 | 3.8 | Globular (Mixed order) |
| 343 | 3.8 | 3.96 | 78805 | 4.4 | Globular (*Mixed order*) |
| +Arginine | | | | | |
| 283 | 2 | 2.56 | 16491 | 1 | Globular (Monomer) |
| 293 | 2 | 2.08 | 16845 | 1.02 | Globular (Monomer) |
| 303 | 2.1 | 2.19 | 17105 | 1.03 | Globular (Monomer) |
| 313 | 2 | 2.47 | 20692 | 1.25 | Globular (*Mixed order*) |
| 323 | 2 | 2.64 | 24186 | 1.4 | Globular (*Mixed order*) |
| 333 | 2.2 | 3.22 | 34953 | 2.11 | Globular (Mixed order) |
| 343 | 2.1 | 3.4 | 43883 | 2.66 | Globular (*Mixed order*) |
| +Sucrose | | | | | |
| 283 | 2 | 2.04 | 7920 | 1 | Globular (Monomer) |
| 293 | 2 | 1.86 | 8202 | 1.03 | Globular (Monomer) |
| 303 | 2 | 1.86 | 8950 | 1.13 | Globular (*Mixed order*) |
| 313 | 2 | 2 | 9533 | 1.2 | Globular (*Mixed order*) |
| 323 | 2 | 2.3 | 11216 | 1.4 | Globular (*Mixed order*) |
| 333 | 2 | 3.1 | 12529 | 1.6 | Globular (*Mixed order*) |
| 343 | 2 | 3.35 | 15234 | 1.9 | Globular (approx. Dimer) |

*The association state of the protein was calculated using $I_0 \alpha$ mass × concentration × time [1,2]

SAXS dataset for dimer was generated by the weighted average of 75% and 25% of the dataset at 313 K and 323 K, respectively.

SAXS dataset for tetramer was generated by weighted averaging 63% and 37% of the dataset at 333 K and 343 K, respectively.

**Supplementary Table S2**

Details of instrumentation and programs used for SAXS processing are tabulated below.

| **Instrument** | **SAXSpace (Anton Paar, Austria)** |
|---|---|
| Collimation | Line Collimation |
| Source | X-rays, CuKα, 0.15414 nm |
| Detector | 1D Mythen |
| Sample to detector distance | 317.06 mm |
| Exposure time & Repeats | One Exposure of 60 minutes for samples & buffer |
| Subtraction from Solutions | Matched Buffer |
| **Programs** | |
| Data collection & Optics Control | SAXSDrive |
| Beam Position Correction | SAXSTreat |
| Buffer Subtraction & Desmearing | SAXSQuant |
| SAXS Intensity File Analysis | ATSAS Suite of Programs v 3.0 |
| **Programs for Shape Restoration** | |
| Guinier Analysis | SAS Data Analysis |
| Distance Distribution Function | SAS Data Analysis |
| Shape Restoration | GASBOR (10 independent runs, no symmetry bias) |
| Superimposition | SUPCOMB [Plugins for PyMol] |
| Theoretical SAS Comparison | CRYSOL |
| **Model Image Generation** | |
| Software | UCSF CHIMERA v 1.14 |

**References**

[1] K. Dhiman, S.K. Nath, Ashish, Monomeric human soluble CD4 dimerizes at physiological temperature: VTSAXS data-based modeling and screening of retardant molecules, J. Biomol. Struct. Dyn. (2020). doi:10.1080/07391102.2020.1771422.

[2] M.D. Badmalia, P. Sharma, S.P.S. Yadav, S. Singh, N. Khatri, R. Garg, Ashish, Bonsai gelsolin survives heat-induced denaturation by forming β-amyloids which leach out functional monomer, Sci. Rep. 8 (2018). doi:10.1038/s41598-018-30951-3.
